# Comprehensive Whole Genome Sequencing Reveals Origins of Mutational Signatures Associated with Aging and Temozolomide Chemotherapy

**DOI:** 10.1101/2024.04.17.590003

**Authors:** Taejoo Hwang, Lukasz Karol Sitko, Ratih Khoirunnisa, Fernanda Navarro Aguad, David M Samuel, Hajoong Park, Banyoon Cheon, Luthfiyyah Mutsnaini, Jaewoong Lee, Shunichi Takeda, Semin Lee, Dmitri Ivanov, Anton Gartner

## Abstract

In a comprehensive study to decipher the multi-layered response to the chemotherapeutic agent temozolomide (TMZ), we analyzed 427 genomes and determined mutational patterns in a collection of ∼40 isogenic DNA repair-deficient human TK6 lymphoblast cell lines. We demonstrate that the spontaneous mutational background is very similar to the aging-associated mutational signature SBS40 and mainly caused by polymerase zeta-mediated translesion synthesis (TLS). *MSH2-/-* mismatch repair knockout in conjunction with additional repair deficiencies uncovers cryptic mutational patterns. We report how distinct mutational signatures are induced by TMZ upon sequential inactivation of DNA repair pathways, mirroring the acquisition of chemotherapy resistance by glioblastomas. The most toxic adduct induced by TMZ, *O^6^*-meG, is directly repaired by the *O^6^*-methylguanine-DNA methyltransferase (MGMT). In *MGMT-/-* cells, mismatch repair (MMR) leads to cell death and limits mutagenesis. MMR deficiency results in TMZ resistance, allowing the accumulation of ∼10^5^ C>T substitutions corresponding to signature SBS11. Under these conditions, N-alkylated bases, processed by base excision repair (BER), limit cell survival. Without BER, 3-meA is read through via error-prone TLS, causing T>A substitutions but not affecting survival. Blocking BER after abasic site formation results in large deletions and TMZ hypersensitization. Our findings reveal potential vulnerabilities of TMZ-resistant tumors.

## Introduction

The genome is under continuous attack from endogenous and exogenous DNA-damaging factors, and genotoxic agents are used for cancer therapy. The lesions that are incurred are either repaired in an error-free manner by the various DNA repair pathways or converted into mutations. The pattern of accumulated mutations reflects the history of genotoxin exposure and the DNA repair deficiencies of a given cell. Mutational signatures classify single nucleotide variants (SNVs) into 96 classes based on the neighboring nucleotides (1). Signatures include information about small insertions/deletions (indels) and structural variants (SVs), such as larger deletions, insertions, inversions, tandem duplications, and translocations. Initially, mutational signatures were computationally derived from cancer genomes based on best-fit approaches and are now compiled in the Catalog of Somatic Mutations in Cancer (COSMIC) (https://cancer.sanger.ac.uk/cosmic), which currently includes 67 single base substitution (SBS) signatures, 23 small indel (ID) signatures, etc. In some instances, the etiology of signatures was easy to infer, e.g., specific mutational patterns were associated with tobacco smoking or exposure to UV light. However, in most cases, their origins remain unclear. Mutational signatures, which are obtained experimentally under controlled conditions by exposing cell lines or organisms to known genotoxins (2), as a result of mutational inactivation of certain DNA repair pathways (3, 4) or as a combination thereof (5), are instrumental in deducing the causes of mutational patterns in cancers but remain underexplored. Recently, studies on human induced pluripotent stem cells (hiPSCs) (3) and haploid HAP1 cells (4) revealed the signatures associated with guanine oxidation (COSMIC signature SBS18 derived from 8-oxoguanine glycosylase OGG1-deficient lines) and mismatch repair deficiency (“universal” MMRd signature RefSig MMR1).

To deduce genotype-specific mutational signatures upon genotoxin exposure, it is necessary to first characterize a spontaneous mutational background of a given cell line. Recent research has identified three universal signatures that, in variable proportions, account for the background somatic mutagenesis in all mammalian species that were investigated and likely reflect three separate mutational processes (6). These signatures are SBS1 (deamination of 5-methylcytosines resulting in C>T substitutions at CpG sites), SBS18 (guanine oxidation leading to C>A substitutions), and SBS5 (featureless signature of unknown etiology). SBS1 and SBS5 are termed clock-like since their mutation counts correlate with age at the cancer diagnosis (7). Remarkably, mutation rates show an inverse correlation with species lifespan, with the final mutation counts at the end of life being very similar across mammals (6). Thus, it appears likely that somatic mutagenesis contributes to cancer and aging. The mechanism leading to the featureless SBS5-like signature and its possible role in aging need to be determined.

To systematically investigate genotoxin-induced mutagenesis across a large panel of cell lines and correlate mutational signatures with cell survival, we focused on temozolomide (TMZ), a chemotherapeutic agent that is used in the treatment of glioblastomas (8). TMZ belongs to S_N_1 (first-order nucleophilic substitution) methylating agents. TMZ treatment gives rise to several methylated bases, which are repaired by different pathways (8, 9). *O^6^*-methylguanine (*O^6^*-meG, 5% of the TMZ-induced adducts (10)) is the most mutagenic and toxic of the lesions due to its ability to pair not only with cytosine but also with thymine. *O^6^*-meG can be repaired by *O^6^*-methylguanine-DNA methyltransferase (MGMT, also known as *O^6^*-alkylguanine-DNA alkyltransferase, AGT), which transfers the methyl group to its cysteine residue in a suicide reaction that restores the guanine base (11). *MGMT* gene is epigenetically silenced in many tumors, thus sensitizing them to TMZ chemotherapy (12). If left unrepaired, *O^6^*-meG is very cytotoxic. Cytotoxicity is mediated by mismatch repair (MMR), and MMR-deficient (MMRd) cells are highly resistant to TMZ. The “futile cycle” model was proposed to explain how MMR of *O^6^*-meG lesions might lead to cell death (13). According to this model, since MMR functions to correct the errors arising during DNA replication, it is not able to excise *O^6^*-meG in the template strand and only repairs a mispaired T in the newly synthesized strand. In the course of filling the daughter strand gap, which resulted from the excision of the mismatched T nucleotide and exonuclease processing, the persisting *O^6^*-meG can again pair with T, thus perpetuating the “futile cycle” of MMR. The ongoing single-strand gap generation apparently interferes with the second round of DNA replication following exposure to TMZ, leading to double-strand (ds) DNA breaks and apoptosis. The observation that cell death ensues only after two rounds of DNA replication (14, 15) is the strongest argument for the “futile cycle” model while an alternative view suggests the “direct signaling” from MMR activation to apoptosis (16). In certain cell types, which are particularly sensitive to replication stress, such as human pluripotent (17) and embryonic stem cells (18), apoptosis is induced already in the first cell cycle following alkylation damage. A recently proposed “repair accident” model postulates that in the quiescent cells, even in the absence of DNA replication, close proximity between the sites of *O^6^*-meG/C mismatch repair and N-alkyl adduct base excision repair might lead to dsDNA breaks, as observed in *N*-methyl-*N*-nitrosourea (MNU)-treated plasmids incubated in *Xenopus* egg extracts (19).

The most common (70%) among the TMZ-induced methylated DNA base adducts is N7-methylguanine (7-meG). 7-meG is neither cytotoxic nor mutagenic. It is, however, recognized by the N-methylpurine DNA glycosylase (MPG, also known as alkyl adenine DNA glycosylase, AAG), which converts 7-meG into an abasic site that is further repaired by the base excision repair (BER) pathway. Overexpression of MPG was shown to increase the sensitivity of cells to methylating agents due to the rapid accumulation of highly toxic abasic sites, which can overwhelm the apurinic/apyrimidinic site endonuclease (APE1)-mediated processing of AP sites (20). N3-methyladenine (3-meA) is a replication-blocking lesion, which is recognized by MPG and repaired by BER (21). Nucleotide excision repair (NER) was implicated as a backup mechanism for the repair of both 7-meG and 3-meA (22), but is thought to only play a minor role. In budding yeast strains deficient in BER, 3-meA can also be repaired by the NER pathway (23) or, alternatively, bypassed by translesion polymerases zeta and REV1 (24). While repair by BER or NER is mostly error-free, translesion synthesis (TLS) might result in nucleotide substitutions either directly at the site of the lesion or at the neighboring stretch of the DNA, which is synthesized by the error-prone polymerases (25).

Although TMZ remains a cornerstone of glioblastoma chemotherapy, in virtually all cases, tumors become resistant to treatments, and the disease remains incurable. It is, therefore, essential to uncover additional vulnerabilities and agents that would prevent the build-up of resistance. Here, we employ an isogenic set of DNA repair gene knockouts to study spontaneous and TMZ-induced mutational patterns. The knockouts were generated using CRISPR in the TK6 TSCER2 B-cell lymphoblastoid cell line, which is near-diploid and is characterized by a stable karyotype and p53 proficiency. Knockouts cover all the major DNA repair pathways (Figure 1A). Similar to a subtype of gliomas susceptible to TMZ treatment, *MGMT* gene in TK6 cells is epigenetically inactivated. In this study, we aimed to determine DNA repair mechanisms that are important for processing TMZ-induced base adducts in the absence of MGMT and MMR with the ultimate goal of finding ways to overcome TMZ resistance. Our results provide a comprehensive analysis of how sequential inactivation of MGMT, MMR, and BER pathways alter the SNV signature from mostly T>C via C>T-dominated COSMIC signature SBS11 to SBS11 with T>A contribution from TLS. These changes likely reflect mutagenesis in glioblastomas before and after TMZ chemotherapy. Our results indicate that N-alkyl adducts are the main killing lesions in the cells that developed tolerance to *O^6^*-meG. We propose XRCC1 and APE1 as potential chemotherapy targets for re-sensitization of tumors that had built up TMZ-resistance in response to treatment.

**Figure 1.**
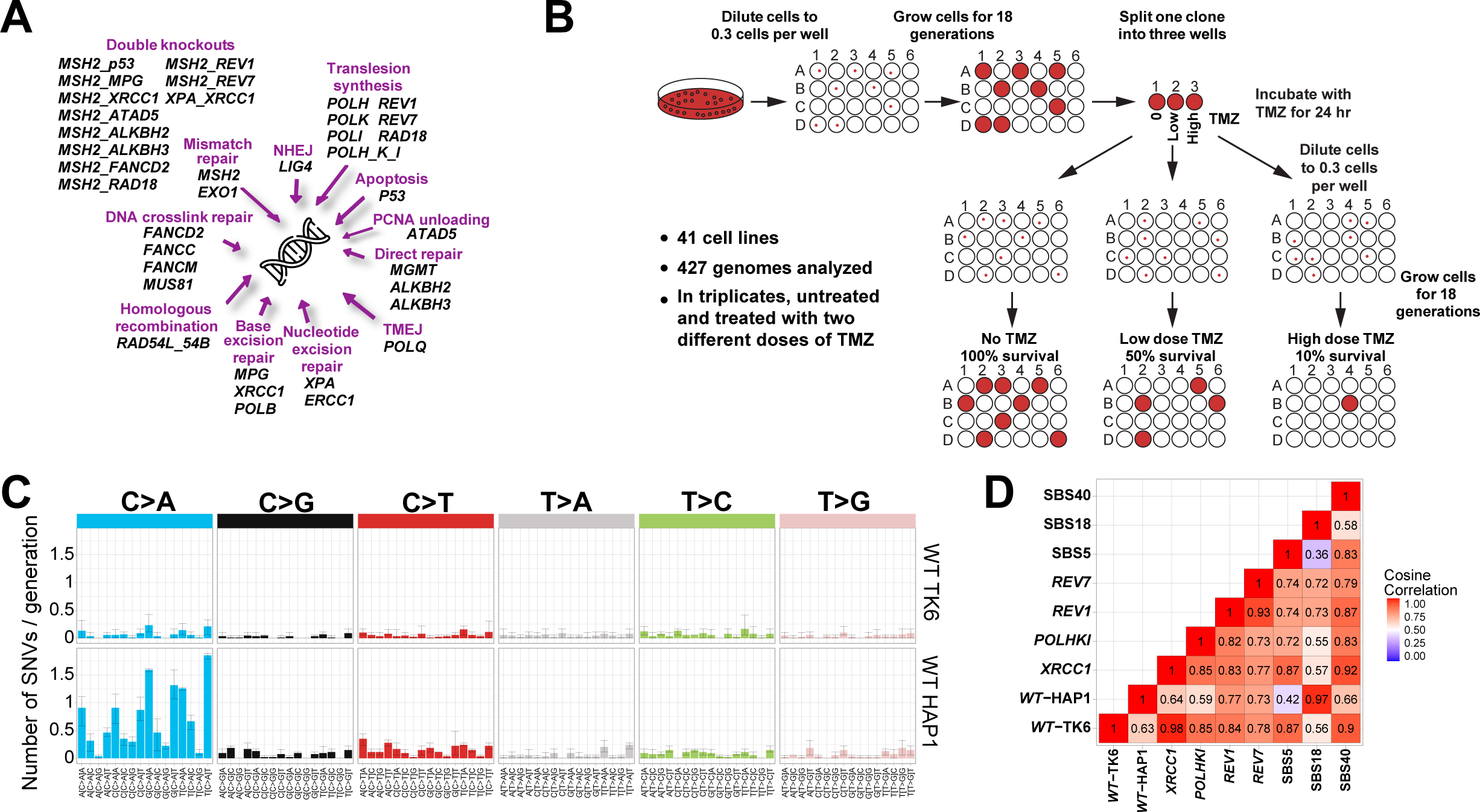
Experimental set-up. **(A)** Collection of DNA repair gene knockouts utilized in this study. **(B)** Experimental workflow. Single cells were obtained by limiting dilution, and corresponding clones were propagated for 18 cell generations to allow mutations to accumulate. Three subclones, each derived from a single cell, were then isolated from the original clonal population and grown for an additional 18 cell generations to amplify the mutations in the founder cell. In the case of TMZ treatment, for each cell line, the parental clone was divided into three wells: one was left untreated, one was treated with the concentration of TMZ causing 50% lethality (LD_50_ ∼2.5 microM TMZ for WT), and one with the concentration of TMZ causing 90% lethality (LD_90_ ∼5 microM TMZ for WT); after 24 hours with TMZ, three subclones were derived from each of the three cell populations. Only sequence variants that differed between the subclones and thus appeared in the course of the experiment were included in the analysis. **(C)** Mutational profile of WT TK6 and HAP1 cells presented in a 96-channel format. Mean ± 95% CI of three subclones. **(D)** Cosine correlations among the spontaneous SNV patterns of the TK6-derived knockout lines and SBS signatures 5, 18 and 40.

## Results

### Spontaneous mutational patterns in TK6 cells

To characterize the contribution of various DNA repair pathways to mutagenesis, we used whole-genome sequencing (WGS) of single-cell progenies to obtain mutation rates and patterns of ∼40 TK6-derived isogenic cell lines with different DNA repair genes inactivated by CRISPR knockout (Figure 1B). Only sequence variants that differed between the subclones and thus appeared in the course of the experiment were included in the analysis. In WT TK6 cells, the spontaneous mutation rate was determined to be approximately 5.4 SNVs, 0.6 indels, and 0.2 SVs per cell generation, which is in good agreement with another TK6 study (26), similar to the rate observed in hiPSCs (27) and significantly lower than the mutation rate in HAP1 cells (4) (Figure 1C). We found that the TK6 cell background mutational pattern closely corresponds to the related signatures SBS5 and SBS40 (28), characterized by an even increase in SNVs (Figure 1D). It was suggested that signature SBS5 could be caused by several mutational processes (29) and might be a combination of several signatures, SBS40 being one of them. The etiology of SBS5 and SBS40 remains unknown. SBS5 and SBS40 are closely related to SBSB, one of the three aging-associated signatures identified in all studied mammalian species (6). Interestingly, the TK6 mutational background is almost entirely devoid of the C>A dominated signature SBS18, which is very prominent in HAP1 (4) (Figures 1C and 1D) and hiPSC cells (3). SBS18 is believed to be caused by oxidative stress, which converts guanines into 8-oxoguanines that can pair with adenines, resulting in C>A substitutions. It was suggested that SBS18 is an *in vitro* signature caused by oxidative stress under cell culture conditions (27, 30). We note that the two cell lines, TK6 (26) and DT40 (31), which do not display SBS18, are lymphoblastic cells growing in suspension. Thus, the oxidative damage resulting in SBS18 may occur due to trypsin treatments of the adherent cells during their passaging.

### TLS polymerase zeta activity is responsible for a background **mutational** signature associated with aging

We found that spontaneous SNVs were decreased by about 1.3X in *REV1-/-* and 2.3X in *REV7-/-* lines (Figure 2A, Supplementary Figure S1A). REV7 is a regulatory subunit of the translesion synthesis (TLS) DNA polymerase zeta, and REV1 plays a role in recruiting polymerase zeta to chromatin, in addition to being a polymerase itself. Polymerase zeta and REV1 were found to be responsible for about half of the spontaneous mutagenesis in budding yeast (32) and in *C. elegans* (33, 34). We suspected that the activity of polymerase zeta might be the underlying cause of background, clock-like and aging signatures SBS5 and SBS40 in human cells. To test this hypothesis, we compared the mutational patterns of the WT TK6 line, *REV1-/-* and *REV7-/-* knockouts, and a triple knockout line deficient in the three Y-family TLS polymerases eta, kappa and iota (*POLH-/- POLK-/- POLI-/-* (35)). We found that REV7 deficiency leads to a dramatic reduction in C>G, T>A, T>C, and T>G counts and results in a mutational spectrum dominated by C>A and C>T substitutions (Figure 2B, Supplementary Figure S1A). Signature SBS40, which is dominant in WT, is not found in the *REV7-/-* spectrum, while SBS18, which is not detectable in WT, accounts for about 32% of mutations in *REV7-/-* (Figure 2C, Supplementary Figure S1B). The mutational pattern of the *REV1-/-* line is intermediate between WT and *REV7-/-* in that both SBS40 and SBS18 can be derived from it. This is consistent with the report that in DT40 cells, ubiquitinated PCNA and REV1 play partially redundant roles in recruiting polymerase zeta to the DNA lesions (36). Curiously, the mutational pattern of a triple knockout *POLH-/- POLK-/- POLI-/-* while overall very similar to WT, has increased numbers of T>A substitutions at C(T>A)T and T(T>A)T (Figure 2B). We conclude that TLS polymerase zeta is responsible for about 50% of spontaneous SNV mutagenesis in TK6 cells. In contrast, TLS polymerases eta, kappa, and iota do not contribute significantly. The remaining 50% of background mutations are probably caused by the oxidation of guanines (SBS18 characterized by C>A substitutions) and possibly deamination of cytosines (C>T substitutions). Interestingly, deamination of 5-methylcytosines, which converts them directly into thymines and causes aging-associated signature SBS1, does not seem to play a significant role in spontaneous mutagenesis in cultured cells since there is no enrichment of C>T substitutions at CpG sequences, which are preferentially methylated. It appears that one of the mechanisms of background mutagenesis in TK6 cells could be spontaneous deamination of cytosines that converts them into uracils, which, if left unrepaired by uracil-DNA glycosylase and BER before DNA replication, result in C>T substitutions.

**Figure 2.**
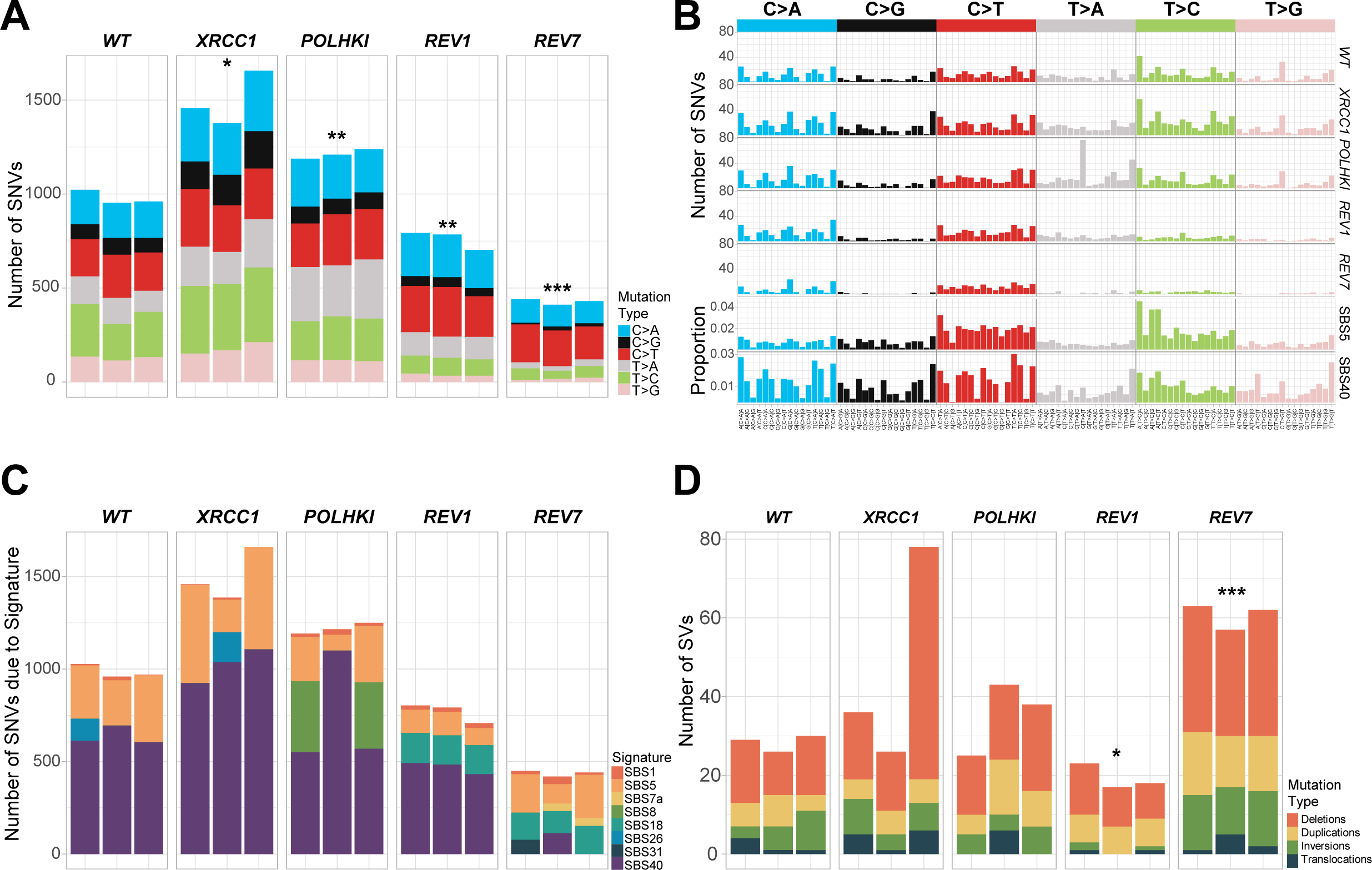
Translesion synthesis by polymerase zeta is a major contributor to spontaneous mutagenesis in TK6 cells. **(A)** Numbers of SNVs accumulated in different cell lines after ∼180 cell generations (10 passages). Significant differences are indicated by * (* p value ≤ 0.05, ** p value ≤ 0.01, *** p value ≤ 0.001, t-test). **(B)** Mutational profiles were obtained in the same experiment as A. (C). Mutational profiles were decomposed into SBS signatures. **(D)** Numbers of SVs from experiment in **A**.

While the SNV rate is decreased in the *REV7-/-* line about 2.3-fold, SVs are increased to a similar degree, especially deletions (Figure 2D, Supplementary Figure S1C). This agrees with a previously reported role of polymerase zeta in protecting the genome from instability (37). Surprisingly, loss of REV1 did not lead to an increase in SVs. Of note, *REV1-/-* but not *REV7-/-* cells display an elevated rate of indels, especially 1 bp deletions (Supplementary Figure S1D), which could be due to the proposed role of REV1 in improving the fidelity of the lesion bypass by polymerase zeta (36). Overall, our results imply that TLS contributes to a flat clock-like signature associated with aging.

We observed a marked increase in spontaneous SNVs in *ERCC1-/-* (p<0.01), *XPA-/- XRCC1-/-* (p<0.01) *EXO1-/-* (p=0.062) and *MSH2-/-* (p<0.01) lines (Figure 3A). In the case of *EXO1-/-*, this increase was below statistical significance due to a high level of variation among the clones. A more moderate but statistically significant increase in SNVs was detected in *POLQ-/-* (p<0.01), *MUS81-/-* (p<0.01), and *POLHKI-/-* triple mutant lines (p<0.01) lines. A trend towards higher SNV levels was also found in *POLB-/-* (p=0.062), *XRCC1-/-* (p=0.05), *RAD54L-/- RAD54B-/-* (p=0.059), *FANCM-/-* (p=0.05), *FANCD2-/-* (p<0.05), and *POLI-/-* (p=0.052) lines (Figure 3A). None of the knockouts, except for *MSH2-/-*, displayed a SNV pattern different from the WT. To obtain higher mutation numbers, we grew the *XRCC1-/-* line for 180 cell generations and compared its mutation pattern to WT grown for the same number of generations. In this experiment, a 1.53-fold increase in the SNV rate in the *XRCC1-/-* line was apparent (p<0.05; 978.7 SNVs in WT vs 1496 SNVs in *XRCC1-/-*), while the SNV pattern remained nearly identical to WT (Fig. 2A-C). XRCC1 serves as a scaffold protein for polymerase beta and ligase III and is essential for the repair of nicks and gaps, including those that are generated in the course of BER. It is conceivable that in the absence of XRCC1, translesion synthesis is augmented, leading to an increased mutational background.

**Figure 3.**
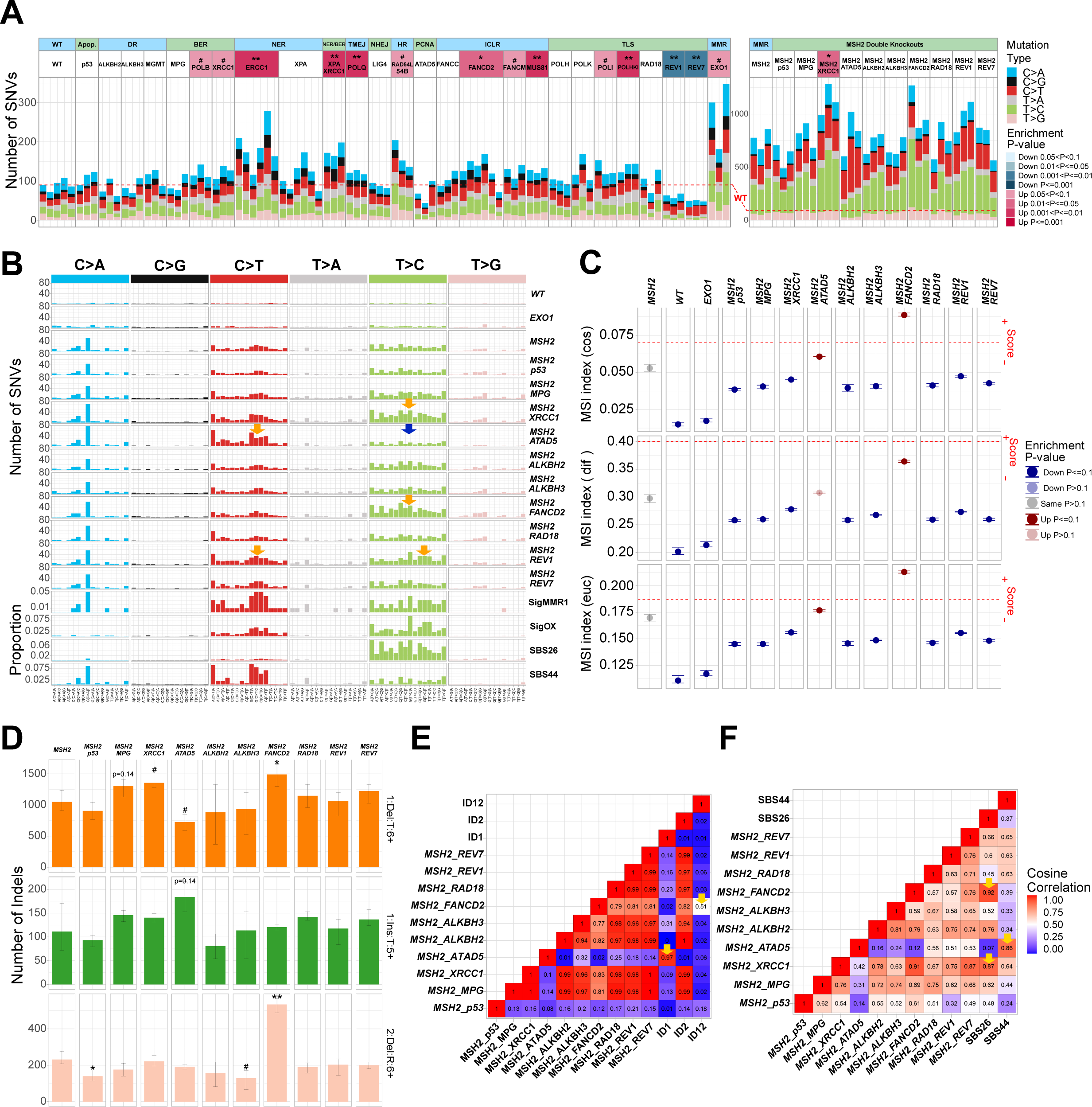
Spontaneous mutagenesis in the collection of TK6 DNA repair mutants. **(A)** Numbers of SNVs accumulated after 18 cell generations in individual subclones. The statistical tests on the left panel are compared to the WT line, and the right panel is compared to the *MSH2-/-* line. **(B)** Mutational profiles in a 96-channel format. Signature SigMMR1 is from (3), Signature SigOX is from (47). **(C)** MSI indexes calculated using MANTIS algorithm: cosine dissimilarity (cos), the stepwise difference (dif), and Euclidean distance (euc). The threshold of instability recommended for the assessment of clinical samples is indicated by a red line. Any value greater or equal to the threshold is considered unstable. Statistical tests are compared to the *MSH2-/-* line. **(D)** Numbers of certain classes of indels accumulated after 18 cell generations: 1 bp deletions in T homopolymers longer than 6 bp (1:Del:T:6+), 1 bp insertion in T homopolymers longer than 5 nt (1:Ins:T:5+), 2 bp deletions in two-nucleotide repeats longer than 6 repeat units (2:Del:R:6+). Mean ± 95% CI of three subclones. Significant differences are indicated by # and * (# p value ≤0.1, * p value ≤ 0.05, ** p value ≤ 0.01, *** p value ≤ 0.001, t-test). **(E)** Cosine correlations among the indel patterns obtained by subtracting *MSH2-/-* indels from double knockouts and indel signatures ID 1, 2, and 12. **(F)** Cosine correlations among the SNV patterns obtained by subtracting *MSH2-/-* SNVs from double knockouts and SBS signatures 26 and 44.

### Mutational signatures in mismatch repair-**deficient** cell lines

*MSH2* knockout leads to a substantial (8.5-fold) increase in SNVs and an even more dramatic (220.5-fold) increase in 1 bp deletions (Supplementary Figure S2A). The distinct SNV pattern of the *MSH2-/-* TK6 line, conforms to the “universal” signature RefSig MMR1 associated with MMR deficiency (MMRd), which was derived from the experimental mutational patterns of different MMRd cell lines (3) (Figure 3B). The RefSig MMR1 signature is effectively a sum of seven signatures, SBS6, 14, 15, 20, 21, 26, and 44, which were computationally derived from MMRd cancers (28), and can be detected in various MMRd cell lines and organisms from yeast to human (38). It is possible that at least some of these seven mutational signatures reflect distinct mechanisms of mutagenesis in MMRd cells. Analysis of double knockouts with *MSH2* proved to be instrumental in revealing replication errors associated with several DNA repair deficiencies, which MMR would likely otherwise correct. We found a change in mutational pattern in the double knockout of *MSH2* and ATPase Family AAA Domain Containing 5 (*ATAD5*) gene, which encodes an ATPase subunit of an alternative replication factor C, whose function is the unloading of the proliferating cell nuclear antigen (PCNA) from chromatin. The C>T substitution count is increased and T>C count decreased in *MSH2-/- ATAD5-/-* compared to *MSH2-/-* (Figure 3B). Therefore, in the SNV spectrum of the *MSH2-/- ATAD5-/-* line, the contribution of computationally derived MMRd-associated signatures is altered with the C>T dominated signature SBS44 increased, and the T>C dominated signature SBS26 decreased (Figure 3F). A likely explanation is that DNA polymerases delta and epsilon have different spectra of errors (39, 40), and their relative contribution to replication is altered in the *MSH2-/- ATAD5-/-* line (Figure S4C). DNA polymerase delta is tethered to chromatin via its interaction with PCNA (41, 42), and DNA polymerase epsilon is recruited via the CMG complex (43, 44). Therefore, it is conceivable that an abnormal accumulation of chromatin-bound PCNA in the absence of ATAD5 will either favor more synthesis or, conversely, create more obstacles for DNA polymerase delta as opposed to epsilon. Another possibility is that the contribution of TLS polymerases to DNA replication is affected by the excess of PCNA in *ATAD5-/-*. A defective interaction between polymerase epsilon and CMG was reported to result in an increased leading strand synthesis by polymerase delta (45) and in enhanced activity of the error-prone polymerase zeta (46).

Conversely, in *MSH2-/- FANCD2-/-* and *MSH2-/- XRCC1-/-* lines, T>C substitutions are increased compared to *MSH2-/-* single knockout. This T>C component is similar to SBS26 and to the SigOX signature, which strongly correlates with oxygen exposure of MMRd cells and was proposed to reflect the effect of oxygen level on polymerase fidelity (47) (Figures 3B and F). Translesion synthesis is likely to be activated in the presence of collapsed replication forks in *FANCD2-/-* or by nicks and gaps accumulating in the XRCC1-*/-* line. No difference in SNV patterns is apparent when comparing WT TK6 with *ATAD5-/-*, *FANCD2-/-* and *XRCC1-/-* lines, probably because of the polymerase error correction by MMR, although the overall SNV count is increased in the *XRCC1-/-* line. A unique pattern was observed in *MSH2-/- REV1-/-* line with both C>T and T>C substitutions somewhat increased and the T>C component different from that in *MSH2-/- FANCD2-/-* and *MSH2-/- XRCC1-/-* lines (Figure 3B).

Microsatellite instability (MSI) is a hallmark of MMRd colon carcinomas (48). However, in some cases, notably in TMZ-treated MMRd gliomas, it cannot be observed by bulk sequencing but only by single-cell sequencing, reflecting the subclonal nature of indels in homopolymers (49). Microsatellite instability (MSI) is not readily observed in cell culture *in vitro*. In HeLa *MSH2* mutants as well as in mouse *MSH2* knockout cells grown in culture for an extended time (80 passages), only limited, so-called “type A” MSI could be detected, but not larger “type B” indels, which are characteristic of MSI-high tumors (50). We could not detect MSI in the *MSH2-/-* TK6 single and double mutant lines when analyzing the MSIplus panel of microsatellite markers, which includes the standard set of 5 loci employed for clinical diagnostics (51) and an additional 13 markers, which are often unstable in tumors (52). However, significant genome-wide MSI could be detected for *MSH2-/-* the *FANCD2-/-* double knockout line using the MANTIS algorithm (53) (Figure 3C). *MSH2-/- FANCD2-/-* MSI index surpasses the threshold using two of three comparison criteria. The MSI score is also increased in *MSH2-/- ATAD5-/-* double knockout line compared to *MSH2-/-* single knockout, although below the recommended threshold. A more detailed analysis of the 83-channel indel pattern revealed that when compared to *MSH2-/-* single knockout, the *MSH2-/- FANCD2-/-* line displays an increase in 2 bp deletions in two-nucleotide repeats longer than 6 repeat units (Figure 3D), which are characteristic of signature ID12 (Figure 3E), as well as an increase in 1 bp deletions (mostly T) at homopolymers, which are characteristic of signature ID2 and are typical of MMR deficiency (28) (Figure 3E). Signature ID2 is attributed to polymerase slippage of the template strand (28). On the contrary, in *MSH2-/- ATAD5-/-* double knockout but not in *MSH2-/-* or *ATAD5-/-* singles, we detected signature ID1, which is attributed to polymerase slippage of the nascent strand and is characterized by 1 bp insertion of T in homopolymers longer than 5 nt (28) (Figures 3D and E). Thus, our results indicate that the mutational inactivation of additional DNA repair pathways enhances microsatellite instability in MMRd cells.

### Mutational patterns in *EXO1-/-* and **Fanconi** Anemia gene knockouts

Interestingly, the knockout of the gene encoding EXO1 exonuclease, which is involved in both HR and MMR, leads to an approximately two-fold increase in the spontaneous SNV count (Figure 3A, although below significance p=0.062 due to clone variability). However, the mutational pattern of *EXO1-/-* is different from *MSH2-/-* (Figure 3B) and instead corresponds to the amplified spontaneous SNV pattern observed in WT TK6 and is similar to SBS5 and SBS40, as was previously observed in *EXO1* knockouts in HAP1 and hiPSCs (3, 4). We also observed a modest increase in 1 bp indels in *EXO1-/-* TK6 line (Supplementary Figure S2A) although not in homopolymer repeats as was reported of *EXO1* knockouts in hiPSCs (3). The MSI score of the *EXO1-/-* line does not differ from the WT TK6 (Figure 3C). Contrary to the previous report (3), we do not observe an increase in double substitutions. There was a minimal increase in tandem duplications and no increase in deletions with microhomology in TK6 *EXO1-/-* (Supplementary Figure S2A and B). These features were reported for HAP1 *EXO1* knockout and interpreted as a sign of HR deficiency (4). Thus, the requirement of EXO1 for HR may differ in different lines.

Deletions with microhomology at breakpoints, characteristic of the HR deficiency-associated signature ID6, are increased in *ERCC1-/-*, *FANCD2-/-*, and *FANCC-/-* lines (Supplementary Figure S2C). These lines show an increase in 1-100 bp deletions and deletions >100 bp, although larger deletions do not exhibit microhomology. Deletions >100 bp are also increased in the *REV7-/-* line (Supplementary Figure S2B). All of these genes function in the Fanconi Anemia pathway. ERCC1 is a regulatory subunit of the XPF nuclease, identified as FANCQ, and REV7 is otherwise known as FANCV. Deletions and other SVs were reported to specifically accumulate in squamous cell carcinomas from Fanconi Anemia (mostly *FANCA*-mutated) patients as opposed to sporadic head-and-neck squamous cell carcinomas (54). Deletions with microhomology, inversions, and tandem duplications were also previously observed in the FANCC-deficient HAP1 line (4). Of interest, the *MUS81-/-* line displayed high levels of spontaneous tandem duplications (Supplementary Figure S2B), as was observed previously in *C. elegans mus-81* mutant (5).

### Mutagenesis induced by the **alkylating** agent temozolomide (TMZ) in WT vs MMR- deficient cells

We determined the sensitivities of our collection of TK6-derived DNA repair gene knockouts to TMZ using clonogenic survival assays (Figure 4A). The TMZ sensitivity varied over a 100-fold range. The most sensitive lines were those deficient in Fanconi Anemia (FA) pathway and ATAD5, and the most resistant lines were deficient for MMR. We then determined the mutational patterns in TMZ-treated clones to investigate how toxicity and mutational signatures are correlated. Cells were exposed to TMZ at concentrations that resulted in 50% and 90% lethality, and TMZ-treated clones were sequenced (Figure 1B). In WT TK6 cells, which do not express MGMT enzyme, TMZ induces mostly C>T substitutions, in agreement with the primary mechanism of TMZ mutagenesis being *O^6^*-meG mispairing with T (Figure 4B). To test that the *MGMT* gene is indeed completely silenced in TK6 cells, we generated a genetic *MGMT-/-* knockout and confirmed that its TMZ sensitivity (Figure 4A) and the TMZ-induced mutational pattern are indistinguishable from the WT (Figure 4B). The number of mutations per microM TMZ was roughly constant across most lines and corresponded to 55.2 C>T substitutions (Figure 5A). This number was increased by approximately 1.4 fold in *RAD18-/-* (82.3 C>T; p<0.001) and *ATAD5-/-* lines (75.5 C>T; p<0.01). Interestingly, in both of these lines, PCNA is dysregulated. RAD18 deficiency largely abolishes PCNA monoubiquitination and reduces TLS (55). However, this is unlikely to be the reason behind an increase in TMZ-induced SNV rate since a triple knockout of TLS polymerases eta, kappa, and iota, as well as REV1 knockout, do not lead to a rise in TMZ-induced C>T substitutions (Figure 5A). Instead, the increased mutagenesis could be due to PCNA dysregulation affecting MMR, the major repair pathway influencing C>T accumulation. Indeed, the number of C>T substitutions per microM TMZ is greatly increased in MMR-deficient *MSH2-/-* (217.8 C>T) and in the *EXO1-/-* line (176.4 C>T), which is expected to be partially MMR-defective. Remarkably, the number of SNVs induced by 500 microM TMZ in *MSH2* knockout reached 100, 000 per cell. This clearly indicates that SNV accumulation is not the cause of lethality after the TMZ treatment of WT cells, in which the SNV count reaches only ∼400 at LD_90_. The proportion of C>T substitutions, which are already dominant among the TMZ-induced mutations in WT TK6 cells, appears to be further increased in the *MSH2-/-* TK6 line (Figure 5A; 77.1% C>T in WT vs 98.3% C>T in *MSH2-/-*). In addition to causing a futile cycle of repair of *O^6^*-meG/T pairs, the MMR pathway may correct some potential C>T substitutions. Alternatively, it is conceivable that TMZ-induced mutations other than C>T arise due to a switch to TLS at the sites where the replication fork encounters an ongoing futile cycle of MMR repair. If this is the case, the proportion of non-C>T SNVs is decreased in the *MSH2-/-* line due to the absence of potential obstacles to replication forks, such as MMR-induced gaps. However, our results from HAP1 cells clearly indicate that MMR reduces the number of C>T mutations (see below).

**Figure 4.**
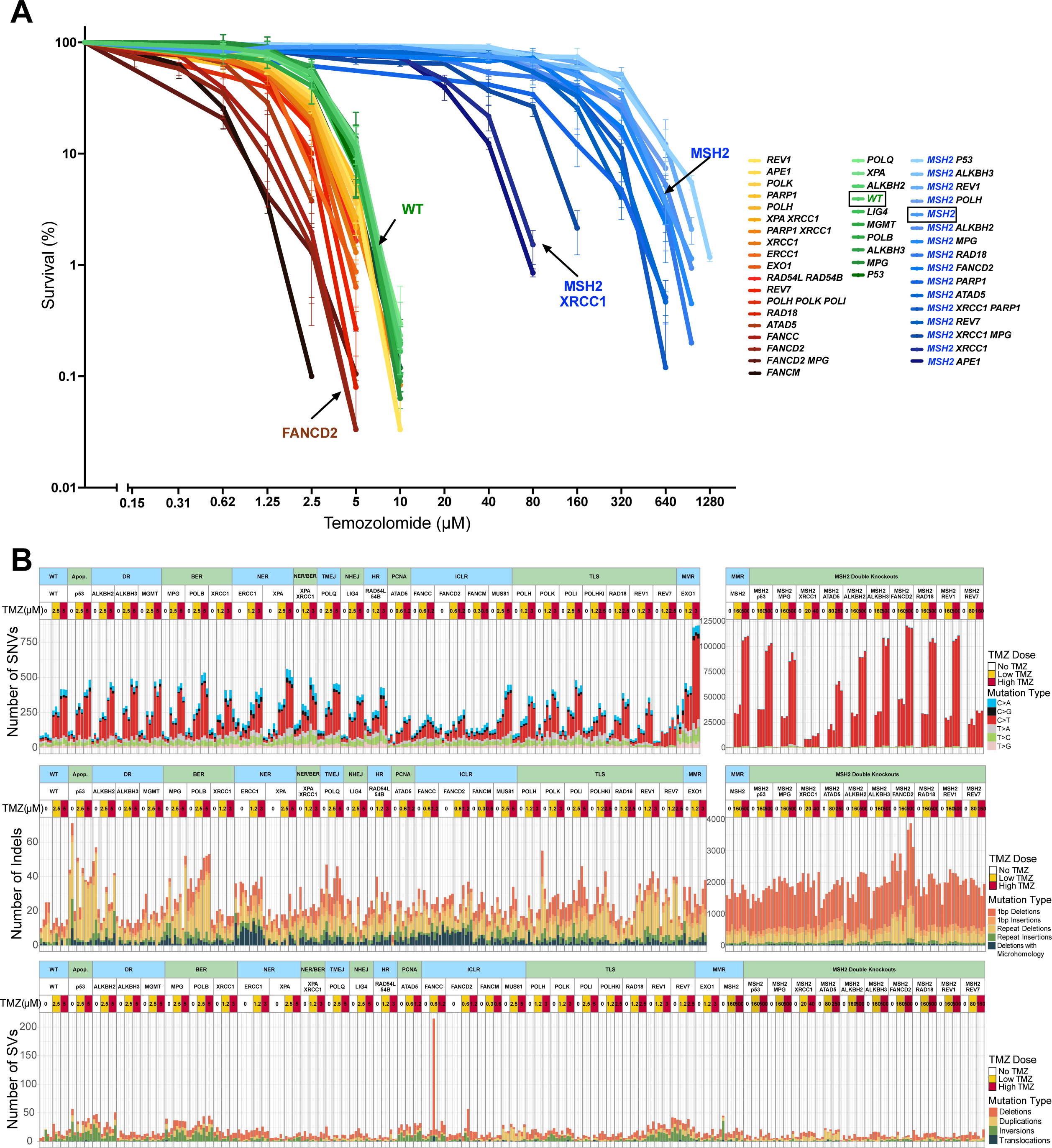
TMZ sensitivities and TMZ-induced mutations in the collection of TK6 DNA repair mutants. **(A)** Cell line sensitivities to TMZ were determined in clonogenic assays. All assays were performed in duplicates. The average survival ± SEM from three independent experiments is plotted using GraphPad Prism 10. (**B)** SNV, Indel, and SV numbers after treatment with low (∼50% survival) and high (∼10% survival) doses of TMZ.

**Figure 5.**
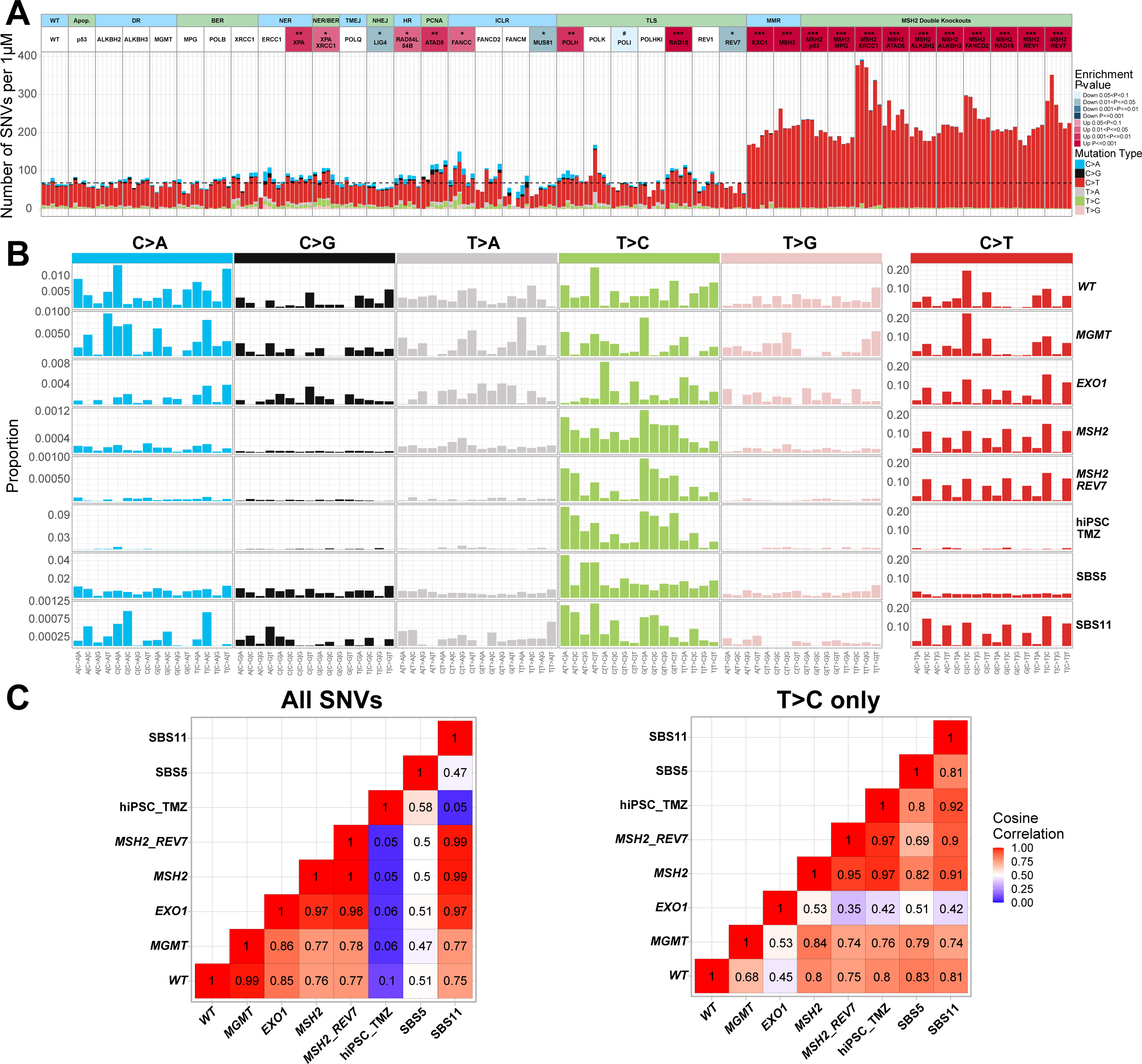
The effect of MGMT and MMR on SNV patterns induced by TMZ. **(A)** Numbers of SNVs induced per microM TMZ. Since different lines were treated with different TMZ concentrations depending on their sensitivity, to compare TMZ-induced mutation rates across the lines, we subtracted the average number of mutations in untreated clones from the average mutation count in TMZ-treated clones and divided the remainder by TMZ concentration. The dashed line corresponds to an average of WT. The statistical tests are compared to the WT line. **(B)** TMZ-induced mutational profiles after subtraction of untreated background. The mutational signature of TMZ-treated hiPSCs is from (2). **(C)** Cosine correlations among the TMZ-induced SNV patterns after subtracting untreated background and SBS signatures 5 and 11. The left panel shows all SNVs, and the right panel shows cosine correlations of T>C patterns only.

The SNV pattern induced by TMZ in *MSH2-/-* line closely resembles signature SBS11 (Figure 5B, Supplementary Figure S3), which was initially derived from TMZ-treated cancers (1) and later demonstrated to result from a combination of TMZ treatment and MMR deficiency selected as a TMZ resistance mechanism (49). Our results confirmed that SBS11 is not observed in WT or any lines other than MMRd after TMZ exposure (Figures 5B and C). Of note, the SNV pattern in the *EXO1-/-* line after the TMZ treatment is very similar to SBS11, even though the *EXO1-/-* spontaneous mutation pattern differs from MMRd lines. Intriguingly, EXO1 deficiency does not lead to TMZ resistance in TK6 cells and instead somewhat hyper-sensitizes to TMZ, which is in disagreement with the result of CRISPR-based genome-wide screen carried out in TMZ-treated MGMT-negative glioblastomas (56). It is conceivable that cell lines differ in their reliance on exonuclease EXO1 activity for MMR as well as HR. Thus, caution should be exercised if signature SBS11 is used as a potential biomarker for MMR deficiency and TMZ resistance, as recently proposed (49).

In addition to C>T dominated SBS11, the TMZ treatment induces an increased rate of background mutagenesis in all MMRd TK6 lines, consistent with an amplified signature SBS5 (Supplementary Figure S3). A similarly amplified background signature was previously reported in TK6 cells after treatment with platinum agents, and it was suggested that spontaneous mutagenesis is accelerated in treated cells (26). Interestingly, knockout of *REV7* in *MSH2-/-* line did not affect the dominant signature SBS11. Still, it reduced the count of the other TMZ-induced mutations (Figures 5B, 7C, and 7D), implying that, like in untreated cells, polymerase zeta activity is responsible for the increase in background mutagenesis following TMZ exposure. Of note, *REV7* knockout did not change the MMRd-specific spontaneous signature in untreated *MSH2-/-* lines (Figure 3B), nor was this signature amplified by TMZ (Figure 5B). In WT TK6 cells, TMZ-induced SNVs other than C>T substitutions were too low to detect any pattern (Figure 5B).

Although C>T substitutions represent the overwhelming majority of SNVs in the TMZ-treated *MSH2-/-* line, TMZ also induced a small number of T>C transitions in this line (Figure 5B). The T>C substitutions are presumably caused by 3-meA and were previously reported to be preferentially induced by TMZ in MGMT-positive hiPSCs (2). Remarkably, the T>C pattern in TK6 *MSH2-/-* cells is very similar to the one in hiPSCs, suggesting that the same mechanism of mutagenesis, most likely TLS, is acting in both lines (Figure 5B and C). T>C substitutions are not observed in TMZ-treated WT TK6 cells, likely because the MMR-dependent futile cycle kills the cells at TMZ concentrations, which are too low to cause a sufficient number of 3-meA lesions.

### TMZ-induced mutagenesis in a MGMT-positive HAP1 line

We wanted to test if C>T-dominated TMZ mutagenesis, which is observed in TK6 cells but not in hiPSCs, is the consequence of TK6 cells being deficient in direct repair of *O^6^*-meG due to epigenetic inactivation of the *MGMT* gene. Thus, we determined the SNV pattern of TMZ-treated WT HAP1 cells, which are MGMT positive, and compared it to the pattern of the HAP1-derived TMZ-treated *MGMT* knockout. *MGMT* knockout HAP1 cells were extremely TMZ-sensitive, with LD_90_ very similar to TK6 cells (Figure 6A). TMZ induced mostly C>T substitutions in these cells (Figure 6B) in a pattern almost identical to WT TK6 (Figure 6E). As expected, WT MGMT-positive HAP1 cells were resistant to TMZ (WT HAP1: LD_50_ ∼150 microM and LD_90_ ∼300 microM vs MGMT-HAP1: LD_50_ ∼3.75 microM and LD_90_ ∼5 microM), and the number of SNVs induced per microM TMZ was greatly diminished compared to *MGMT* knockout HAP1 (Figure 6D). Surprisingly, the six subclones of the same parental WT HAP1 clone, which were treated with TMZ and sequenced, differed greatly in the number and type of mutations they accumulated. Three subclones accumulated 1, 100-6, 400 mutations, mainly C>T substitutions, similar to TK6. Two accumulated only ∼700 mutations, mostly T>C, and one accumulated both C>T and T>C at a similar ratio (Figure 6B). Intriguingly, the T>C pattern in the two clones with the lowest mutation count was very similar to the one observed in the TMZ-treated hiPSC cells (2), and the C>T pattern in the three clones with the highest mutation count was similar to signature SBS11 of the TMZ-treated MMRd lines (Figure 6E). We suggest that MGMT expression in HAP1 cells is variable, with MGMT-low subclones spontaneously appearing at high frequency, probably due to a partial epigenetic inactivation of the *MGMT* gene. Indeed, when we performed a Western blot to examine MGMT protein expression in six HAP1 subclones derived from the same clone without drug treatment, we observed an extremely low level of MGMT protein in two of them (Figure 6C). Of note, the subclones with high C>T levels following the TMZ exposure cannot be MGMT-null since they survived treatment with TMZ concentrations 60 times higher than LD_90_ for *MGMT* knockout HAP1. However, the MGMT-low subclones are less efficient in directly repairing *O^6^*-meG and, therefore, accumulate C>T substitutions. The similarity of their SNV pattern to SBS11 suggests that MMR capacity is overwhelmed by the huge numbers of *O^6^*-meG mispairing with T (Figure 6E). Thus, signature SBS11 is indicative of *O^6^*-meG/T pairs being converted into C>T substitutions without MMR correction and might be observed in the cell lines, which have been treated with high TMZ concentrations (in the 100 microM range), even if they are not MMR deficient.

**Figure 6.**
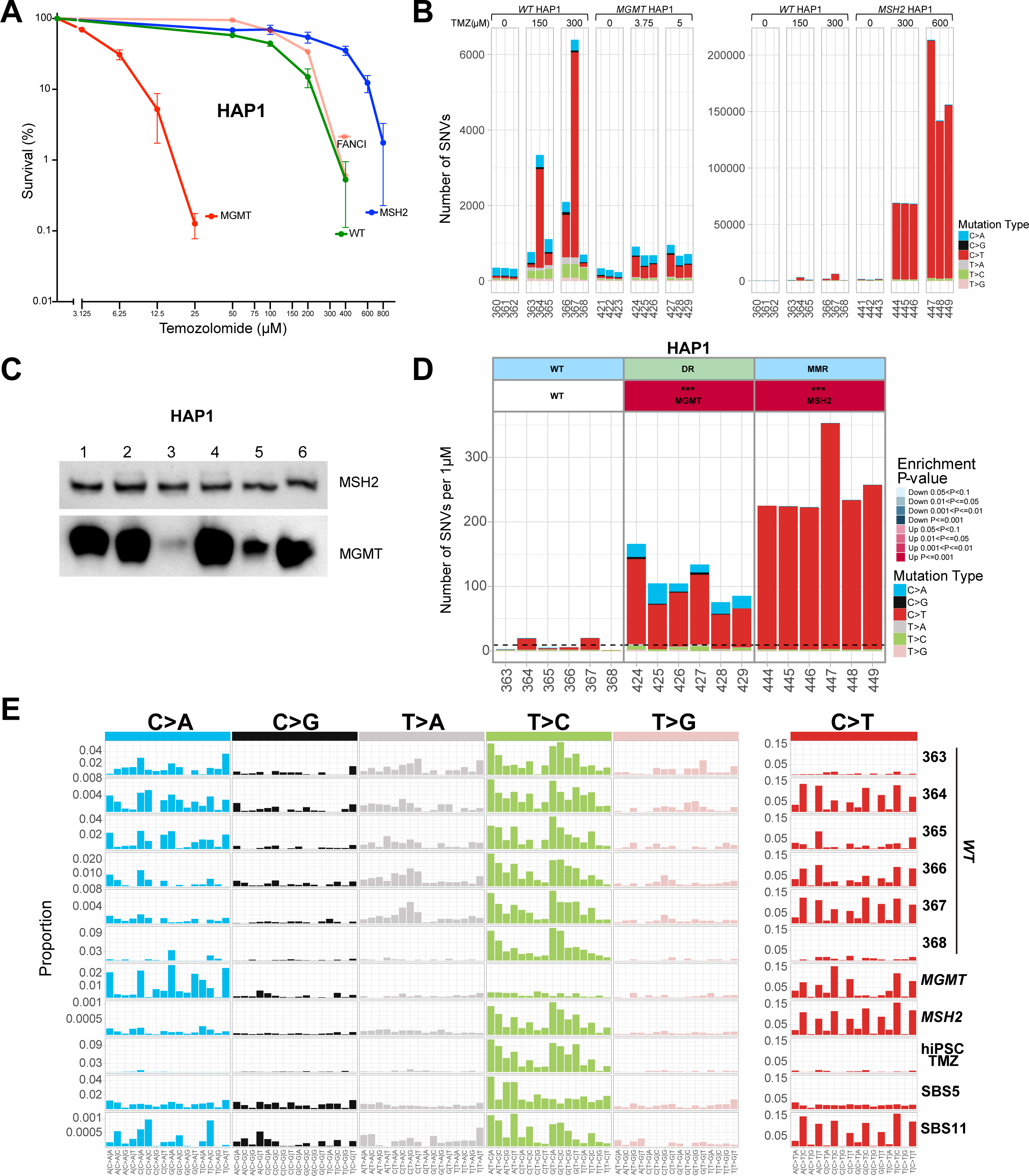
TMZ-induced mutagenesis in HAP1-derived lines. **(A)** HAP1-derived cell line sensitivities to TMZ were determined in clonogenic assays. All assays were performed in duplicates. The average survival ± SEM from three independent experiments is plotted using GraphPad Prism 10. **(B)** Numbers of SNVs in HAP1 subclones after treatment with the indicated TMZ doses. **(C)** Western blot showing different expression levels of the MGMT enzyme in a set of WT HAP1 subclones without TMZ treatment. These subclones were obtained from a different experiment than in d. **(D)** Numbers of SNVs induced per microM TMZ in HAP1 subclones after subtraction of untreated background. The dashed line corresponds to an average of WT. **(E)** TMZ-induced mutational profiles in HAP1 subclones after subtraction of untreated background. The signature of TMZ-treated hiPSCs is from (2). Panels **B, D,** and **E** are from the same experiment. Subclone ID numbers are indicated under the graphs.

Disruption of *MSH2* in WT HAP1 cells rendered them even more resistant to TMZ (LD_50_ ∼300 microM and LD_90_ ∼600 microM). Interestingly, when WT and *MSH2* knockout HAP1 cells are treated with the same TMZ concentration of 300 microM, *MSH2* knockout cells accumulate ∼10 times more substitutions than WT clones with high mutation numbers; ∼ 70, 000 SNVs per cell, mostly C>T (Figure 6B), indicating that MMR prevents the majority of C>T mutations. The number of C>T substitutions per cell is even higher, up to 200, 000 in HAP1 *MSH2* knockout treated with 600 microM TMZ. The SNV pattern of TMZ-treated MSH2-deficient HAP1 cells closely resembles signature SBS11. Thus, this signature can be observed in both MGMT-positive and negative MMRd cells after TMZ treatment.

### Base excision repair deficiency re-sensitizes MMR-**deficient** cells to TMZ

To understand which DNA repair deficiencies might re-sensitize *MSH2-/-* cells to TMZ, we created a set of double knockouts based on the *MSH2-/-* TK6 line. We reasoned that if *O^6^*-meG, despite remaining mutagenic, is rendered non-toxic in MMRd cells, other TMZ-induced DNA base adducts will become limiting for cell survival, especially 3-meA, which acts as a replication blocker. Therefore, we tested if the inactivation of BER, direct repair of 1-meA and 3-meC, TLS, or FA pathways sensitize *MSH2-/-* cells to TMZ (Figures 4A and 7A). One double knockout line, *MSH2-/- TP53-/-*, was somewhat more resistant to TMZ, indicating that at least a fraction of TMZ-induced cell deaths might be attributed to p53-dependent apoptosis (Figure 4A). *MSH2-/- ATAD5-/-*, *MSH2-/- REV7-/-* and *MSH2-/- PARP1-/-* lines were moderately more sensitive to TMZ, while the *MSH2-/- XRCC1-/-* line was dramatically more sensitive to TMZ than *MSH2-/-* (LD_50_ ∼20 microM and LD_90_ ∼40 microM) (Figures 4A and 7A). Sequencing revealed that TMZ induced deletions larger than 200 bp, specifically in the *MSH2-/- XRCC1-/-* line (Figure 7B). These deletions likely arise from dsDNA breaks induced by unrepaired nicks that accumulate without XRCC1. XRCC1 is involved in BER, and mutations in the *XRCC1* gene are known to cause extreme sensitivity to alkylating agents, such as EMS and MMS (57). However, in our experiments, *XRCC1* knockout in TK6 WT caused only a minor increase in TMZ sensitivity, indicating that it does not play a role in repairing *O^6^*-meG lesions. On the contrary, *XRCC1* knockout in the *MSH2-/-* line resulted in a major hypersensitization to TMZ (Figure 7A). This could be due to the two non-mutually exclusive mechanisms. In *MSH2-/-* line treated with a high TMZ concentration, 7-meG and 3-meA lesions engage BER, which requires XRCC1 to function as a scaffold for polymerase beta and ligase III required for the last step of filling the gaps and sealing the nicks. Alternatively, it was proposed that XRCC1’s role is to suppress the excessive PARP1 trapping on SSB intermediates, which, when left unchecked, might lead to NAD+ depletion and cell death (58, 59). To test these hypotheses, we generated triple knockouts. Inactivation of the BER pathway in *MSH2-/- XRCC1-/- MPG-/-* line partially reversed *MSH2-/- XRCC1-/-* hypersensitivity to TMZ, indicating that the inability to complete BER repair accounts for some but not all TMZ sensitivity of *MSH2-/- XRCC1-/-* line (Figure 7A). Remarkably, *MSH2-/- XRCC1-/- PARP1-/-* cells are even more resistant to TMZ than *MSH2-/- XRCC1-/- MPG-/-*, suggesting that after TMZ treatment, PARP1 trapping might occur not only during BER but also at stalled replication forks (60) or unligated Okazaki fragments (61) and XRCC1 might be required for PARP1 release during different DNA repair processes. To confirm that blocking the BER pathway at an intermediate step re-sensitizes MMRd cell lines to TMZ, we generated *APE1-/-* single and *MSH2-/- APE1-/-* double knockouts. *APE1* encodes apurinic/apyrimidinic endonuclease, which performs the second step in the BER pathway. Like XRCC1, APE1 deficiency had a minimal effect on the WT TK6 cells’ sensitivity to TMZ but dramatically sensitized the MSH2-*/-* line (Figure 7A).

**Figure 7.**
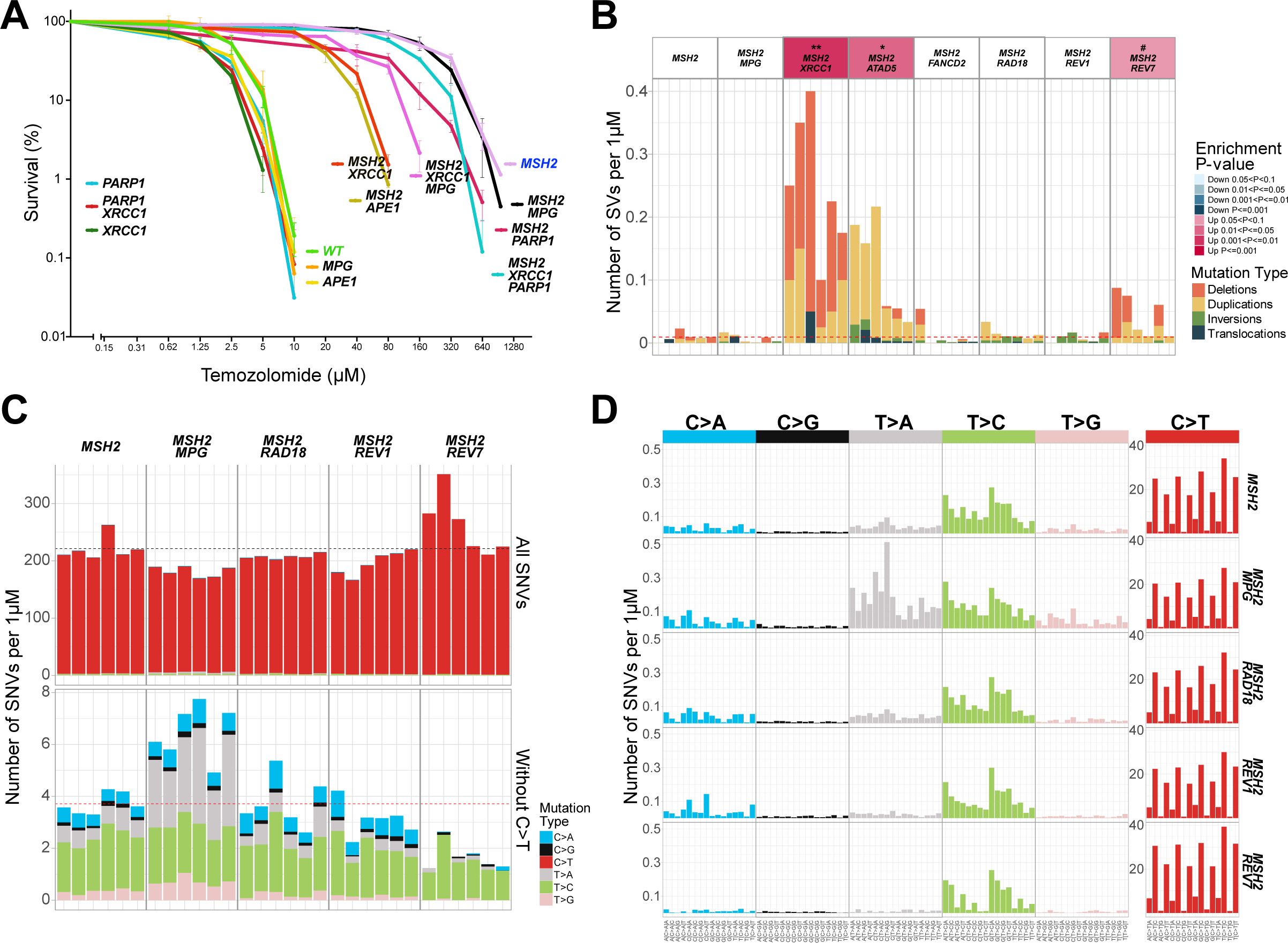
The role of BER and TLS in the survival of MMRd lines after treatment with high TMZ doses. **(A)** Cell line sensitivities to TMZ were determined in clonogenic assays. All assays were performed in duplicates. The average survival ± SEM from three independent experiments is plotted using GraphPad Prism 10. **(B)** Numbers of SVs induced per microM TMZ after subtracting untreated background. The dashed line corresponds to an average of WT. **(C)** Numbers of SNVs induced in *MSH2-/-* double knockouts per microM TMZ after subtraction of untreated background. The dashed line corresponds to an average of *MSH2-/-*. **(D)** TMZ-induced mutational profiles in *MSH2-/-* double knockouts per microM TMZ after subtraction of untreated background.

Knockout of the MPG gene, which encodes N-methylpurine DNA glycosylase, the enzyme catalyzing the first step in BER by removing the alkylated base and creating the abasic site, did not result in TMZ hypersensitivity either in WT or in *MSH2-/-* lines (Figure 7A). However, the spectrum of TMZ-induced mutations in *MSH2-/- MPG-/-* line changed compared to *MSH2-/-*. In *MSH2-/- MPG-/-* T>A substitutions were significantly increased (Figures 7C, 7D, and Supplementary Figure S4A). The reason is likely that in the absence of BER, which efficiently repairs 3-meA error-free, the replication block imposed by 3-meA is relieved by error-prone TLS, leading to the accumulation of T>A substitutions. Consistent with this view, knockout of *RAD18* affected neither TMZ resistance (Figure 4A) nor mutagenesis (Figures 7C, 7D, and Supplementary Figure S4A) in the *MSH2-/-* line. Thus, BER and TLS likely act redundantly to overcome the harmful effects of 3-meA on cell viability.

### Fanconi Anemia and *ATAD5* knockouts hyper-**sensitize** TK6 cells to TMZ exposure

The most TMZ-sensitive lines in our collection are *FANCM-/-* (LD_50_ ∼0.312 microM and LD_90_ ∼0.625 microM), *FANCC-/-*, *FANCD2-/-* and *ATAD5-/-* (Figure 4A). This is unexpected since neither of these genes is known to be involved in repairing alkylation damage. Fanconi Anemia (FA) DNA repair pathway’s primary function is believed to be the repair of DNA interstrand crosslinks, although it is also involved in the protection of stalled replication forks (62). TMZ hypersensitivity of glioblastoma cells deficient in the FA pathway was observed in the CRISPR-based genome-wide screen (56). In addition, a different CRISPR screen revealed that disruption of FA genes hypersensitized RPE1 cells to alkylating agents MMS and MNNG (63). However, the mechanism of how the FA pathway is involved in resistance to alkylating agents remains to be investigated. One possibility to consider is that 7-meG, the most abundant among the TMZ-induced adducts, triggers BER and abasic sites generated by N-methylpurine DNA glycosylase (MPG) spontaneously convert into interstrand crosslinks, thus necessitating their repair by the FA pathway. We could rule out this mechanism since the double knockout *MPG-/- FANCD2-/-* is as sensitive to TMZ as *FANCD2-/-* (Figure 4A). Alternatively, the FA pathway may be required to overcome the toxicity of *O^6^*-meG. Since the toxicity of *O^6^*-meG depends on MMR, we constructed an *MSH2-/- FANCD2-/-* double knockout, which was found to be as TMZ-resistant as *MSH2-/-* (Figure 4A). Thus, most likely, the FA pathway is involved in the protection of replication forks encountering sites of an ongoing futile cycle of MMR repair where *O^6^*-meG is paired with T. Sequencing revealed that TMZ-induced deletions of all sizes in FA knockouts (Supplementary Figure S4B), although in case of *FANCM-/-* below statistical significance, probably due to very low concentration of TMZ that this line can tolerate. Remarkably, the NER-defective *ERCC1-/-* line also displays an increased rate of spontaneous deletions (Supplementary Figure S2), similar to FA ko. However, in this line, deletions are not induced dramatically by TMZ, unlike what we observe in FA-deficient lines (Supplementary Figure S4B). Consistent with the observed correlation between deletion induction and TMZ sensitivity, the *ERCC1-/-* line is only modestly hypersensitive to TMZ (Figure 4A). Likewise, TMZ, even at 500 microM concentration, did not induce deletions in *MSH2-/- FANCD2-/-* double knockout (Figures 4B and 7B). In the HAP1 line, *O^6^*-meG is directly repaired by MGMT, rendering it resistant to TMZ. In this line, FANCI knockout did not induce TMZ hypersensitivity, consistent with what was observed in an MGMT-positive subset of glioblastomas (56) (Figure 6A). We conclude that the FA pathway is required to overcome MMR- dependent toxicity of *O^6^*-meG. Alternatively, the FA pathway may promote cell survival by attenuating MMR activity, which is associated with break induction and ensuing apoptosis. The antagonistic relationship between these two repair pathways has been previously suggested (64), although the mechanism remains to be elucidated.

Since the futile cycle of MMR repair of *O^6^*-meG paired with T is the primary mechanism of TMZ toxicity, overstimulation of MMR might be the reason why ATAD5 deficiency hypersensitizes to TMZ. The main function of ATAD5 is to unload PCNA from chromatin. As PCNA plays an important role in MMR, excessive PCNA on DNA may lead to overactivation of the MMR pathway and increased futile cycles. Indeed, ATAD5 (Elg1) deficiency in budding yeast results in over-recruitment of MutS and accumulation of MutH foci (65). Similarly, in human cells, downregulation of ATAD5 leads to MSH2 accumulation on chromatin (66) (Supplementary Figure S4C). However, this might not be the only mechanism involved since the *MSH2-/- ATAD5-/-* double knockout line, while much more TMZ-resistant than *ATAD5-/-*, is still somewhat more sensitive to TMZ than *MSH2-/-*. Curiously, TMZ induces tandem duplications in *MSH2-/- ATAD5-/-* line (Figure 7B), typical of homologous recombination (e.g., BRCA1) deficiency (67). However, the size of TMZ-induced tandem duplications is uncharacteristically small (median ∼400 bp vs ∼10 kb in breast cancers (68, 69)). In addition, TMZ treatment results in indels, including deletions flanked by microhomology in *MSH2-/- ATAD5-/-* (Supplementary Figure S4D), another indicator of HR deficiency (28). ATAD5 was recently shown to be involved in HR by promoting short-range resection (70). The HR requirement for TMZ resistance might be increased when *MSH2-/-* line is treated with very high TMZ concentration and the replication-blocking adduct 3-meA accumulates. This would explain why tandem duplications are not observed in the *ATAD5-/-* line treated with low TMZ.

Interestingly, TMZ induces dinucleotide deletions in poly A repeats longer than 12 nt in the POLB-*/-* line deficient in DNA polymerase beta, which is involved in the final step of BER (Supplementary Figure S4E). Persistent nicks generated due to incomplete BER of 3-meA may lead to such repeat instability.

### *ALKBH2-/-* and *ALKBH3-/-* knockouts are not hypersensitive to TMZ

In addition to *O^6^*-meG, 7-meG, and 3-meA, TMZ induces two more base adducts, N1- methyladenine (1-meA) and N3-methylcytosine (3-meC), which interfere with the DNA base pairing and thus are weakly mutagenic but highly cytotoxic since they can disturb DNA replication. Human alpha-ketoglutarate-dependent dioxygenase AlkB homolog 2 (ALKBH2) directly reverses these lesions and was reported to contribute to glioblastoma resistance to TMZ (71). Another ALKB homolog, ALKBH3, demethylates 1-meA and 3-meC in single-stranded (ss) DNA. It forms a complex with ASCC helicase, which unwinds dsDNA, thus enabling its activity (72) ALKBH3 is required for resistance to alkylating agents in some cancer cell lines but not in others (72) and might act redundantly with ALKBH2 (72, 73). In our study, knockout of neither *ALKBH2* nor *ALKBH3* resulted in TMZ hypersensitization in WT or *MSH2-/-* TK6 cells (Figure 4A).

All in all, we provide a comprehensive view of how mutational signatures are molded by multiple DNA repair pathways. Our approach is relevant for understanding how chemotherapeutic agents act and reveals a new vulnerability of TMZ-resistant cells with important implications for glioblastoma treatment.

## Discussion

In this study, we focused on genotoxin-induced mutations using WGS of an isogenic set of human DNA repair gene knockout lines. As a first step, we characterized spontaneous mutation rates and patterns in the various repair-defective lines. A compelling result of these initial experiments is that TLS polymerase zeta deficiency conferred by the knockout of its regulatory subunit REV7 leads to a reduced SNV rate and largely eliminates signature SBS40 from the mutational background.

SBS40 is related to a clock-like signature SBS5, which becomes more pronounced as organisms age (7) and is similar to SBSB, one of the three universal aging-associated signatures identified in mammals (6). The gradual accumulation of mutations over the organism’s lifespan (6), along with the epigenetic changes (74, 75), was long suspected to be the underlying cause of senescence, although it remains to be established if translesion-dependent SNV accumulation merely correlates with aging or has a causative role. Polymerase zeta function presents a trade-off: it protects the genome from potentially much more harmful deletions by increasing the rate of SNVs. While preparing this manuscript for publication, a paper by Szuts’ lab reported, using RPE-1 cells, that *REV3L* and *REV1* knockouts displayed a reduced rate of SNV accumulation, with signature SBS40 eliminated from the mutational spectra (76). As noted by the authors, a high proportion of cell culture-induced oxidative signature SBS18 in the RPE-1 mutational background complicates using this cell line as a model to study spontaneous mutagenesis of human cells. A low SNV rate and SBS40-dominated spontaneous spectra of TK6 cells, employed in our study, better recapitulate human cell mutagenesis *in vivo*. In TK6, the C>A dominated oxidative signature becomes unmasked only when the “flat” signature SBS40 is reduced in *REV7-/-* knockout.

Whereas the error-prone translesion synthesis contributes to spontaneous SNV mutagenesis, the mismatch repair pathway corrects the replication errors and limits the nucleotide substitution rate. MMR inactivation results in a dramatically elevated mutagenesis with a distinct mutational pattern, which is observed in various tumors, cell lines, and organisms from yeast to humans (38). Correction of mismatches prevents mutations caused by various DNA repair gene deficiencies, and double knockouts with MMR genes, e.g., *MSH2-/-*, help to unmask the new, primary mutational patterns. Intriguingly, the initial seven signatures, which were computationally derived from MMRd colon cancer, and then appeared to “merge” in the experimentally obtained “universal” signature RefSig MMR1, might represent distinct and separable mutational mechanisms. For example, our results demonstrate that C> T-dominated SBS44 and T> C-dominated SBS26 are up- and down-regulated in an opposing manner in *MSH2-/- ATAD5-/-* and *MSH2-/- FANCD2-/-* double knockouts. The exact mechanisms that cause these signatures remain to be identified.

Having characterized spontaneous mutagenesis, we comprehensively analyzed the mutational patterns associated with genotoxin-induced lesions, using the clinically relevant methylating agent TMZ as a model drug. The strength of our approach is combining clonogenic cytotoxicity assays with WGS to reveal mutational patterns. Cell survival following the drug treatment does not always correlate with the mutational burden. We employed whole genome sequencing to gain deeper insight into how TMZ kills the cells. WGS revealed that TMZ-resistant *MSH2-/-* cells treated with a very high drug concentration accumulate up to a hundred thousand SNVs per cell without losing viability. On the other hand, induction of a relatively small number of deletions in TMZ-treated FAd and *MSH2-/- XRCC1-/-* lines serves as an indicator of dsDNA breaks and correlates with cell death. Deletions and other SVs seem to be linked to cell lethality and, therefore, are challenging to detect in the few clones that manage to survive the treatment.

One of our study’s main conclusions is that in order to deduce the genotoxin-induced mutational patterns, which reveal the contribution of various repair pathways, these pathways need to be inactivated in a stepwise manner (Figure 8A). Although reduced repair gene functionality is specifically selected for in tumors, the associated signatures do not become apparent in most cases, likely because a moderate increase in lifetime mutational burden leads to an exponential increase in cancer risk (5, 77).

**Figure 8.**
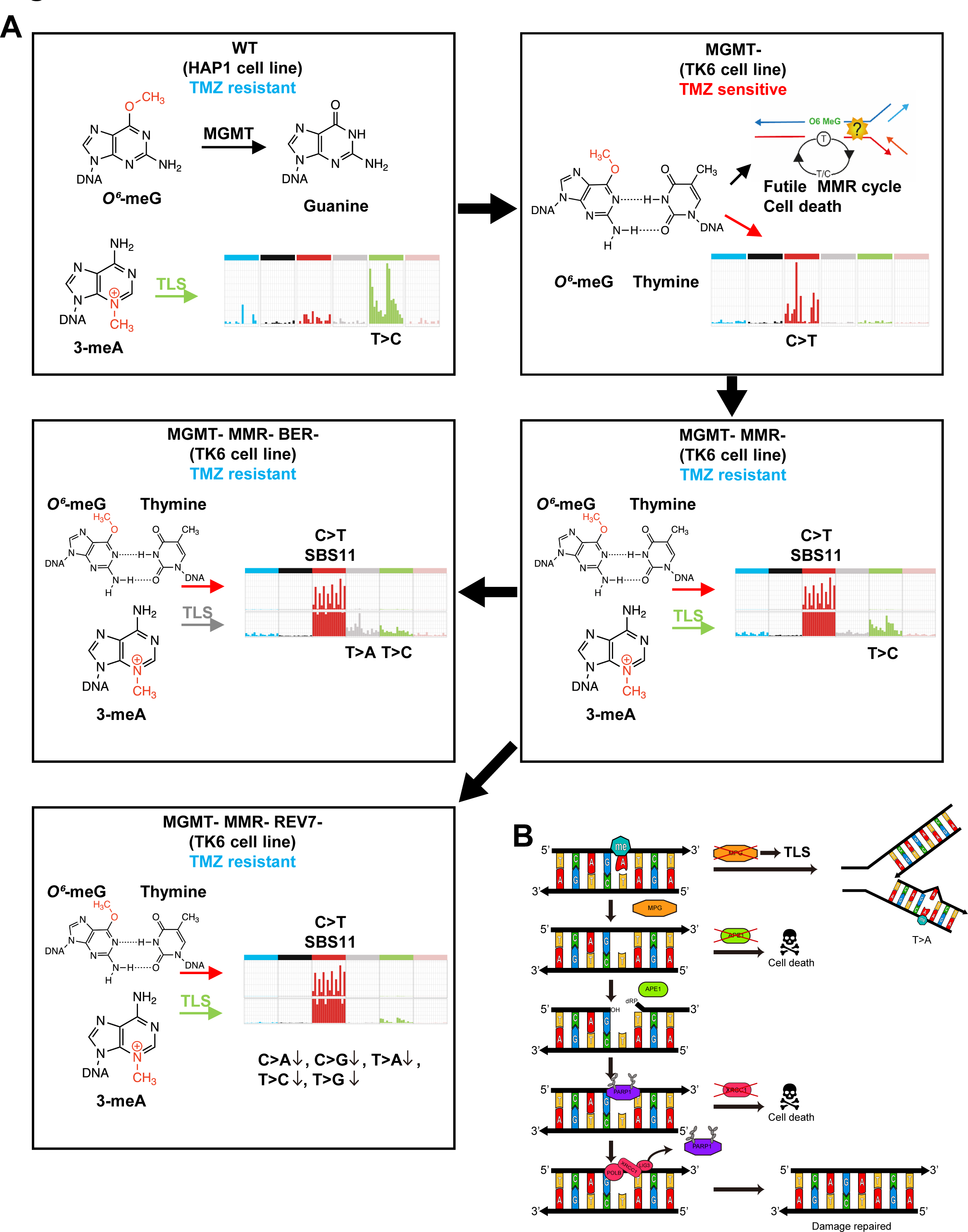
Model of the multi-layered response to TMZ. **(A)** Model of how the TMZ-induced mutational signatures change upon inactivation of different DNA repair pathways (See Discussion for details). **(B)** BER and TLS mediate resistance to 3-meA.

Our study explains the results of a previous comprehensive analysis of mutational signatures, which were induced in the wild-type hiPSCs by a set of diverse genotoxic agents (2). In particular, treatment with MMS and MNNG methylating agents at lethal concentrations triggering DNA damage response, resulted in no distinct mutational signatures in hiPSCs, while TMZ induced only a small number of T>C substitutions, very different from C>T dominated SBS11 observed in TMZ-treated cancer samples (2). The T>C changes are consistent with the mutational pattern we observed in some clones of the HAP1 cell line, which presumably express high levels of the MGMT enzyme that directly repairs *O^6^*-meG, the most toxic of TMZ-induced base adduct in MMR-proficient cells. T>C mutagenesis is likely mediated by TLS through replication-blocking 3-meA lesions. However, the TMZ-induced T>C signature is not observed in MGMT-negative tumors, which are sensitive to, and therefore selected for, TMZ treatment. MGMT deficiency allows *O^6^*-meG accumulation, which drives C>T substitutions. However, due to MMR-mediated futile repair cycles, *O^6^*-meG adducts ultimately become converted to lethal lesions, presumably dsDNA breaks, leading to cell death and limiting mutation accumulation. Inactivation of the MMR pathway results in resistance to very high TMZ concentrations and massive numbers of C>T substitutions conforming to signature SBS11, first derived from the genomes of TMZ-treated tumors.

Signature SBS11 was first computationally derived from the TMZ-treated tumors (1) and then experimentally shown to result from a combination of the TMZ treatment and MMR deficiency (49). Here, we report a further insight into the origin of SBS11. This signature is observed in some clones of MGMT+ MMR+ cells (presumably those with a lower level of MGMT expression), as well as in MGMT+ MMRd and MGMT- MMRd cells, which were treated with high concentrations of TMZ (in the 100 microM range). However, it is also displayed by the MGMT-negative *EXO1-/-* cells treated with as low as 3 microM TMZ but not in MGMT- WT TK6 cells or other mutants. SBS11 is characterized by a high rate of C>T substitutions in CC and CT but not in CA and CG sequences. This pattern appears to be intrinsic to TMZ mutagenesis in the absence of MMR and likely reflects a neighboring base bias in the methylation of guanines or in the propensity of *O^6^*-meG to mispair with T. It is worth noting that the product of TMZ hydrolysis, 5-(3-methyl 1-triazenyl) imidazole-4-carboxamide (MTIC), was previously reported to preferentially methylate the N7 positions of the inner guanines in the tracks of 3-4 Gs rather than individual Gs (78). If the same sequence preference holds for *O^6^*-meG, it will correspond to C>T substitutions in CCC sequences. The C>T pattern is modified by MMR as reflected in the

TMZ-induced signature from the WT cells, where there are fewer C>T substitutions in GC sequences and more C>T substitutions in CCC and CCT. It is possible that *O^6^*-meG/T pairs, followed by a C, are preferentially corrected. In summary, SBS11 is observed in TMZ-treated cells, where MMR is either deficient or overwhelmed by the large numbers of *O^6^*-meG adducts mispairing with T. This signature is expected to be also observed in tumors, which acquired TMZ-resistance due to the mechanisms other than the MMR deficiency if they were exposed to high TMZ concentrations and the MMR capacity was exceeded.

Since *O^6^*-meG is not toxic in the absence of MMR, TMZ cytotoxicity in MMRd lines is mediated by the other base adducts, especially 3-meA. Mutagenesis via 3-meA in MMRd cells is largely prevented by the error-free BER pathway and results in a relatively modest number of T>C substitutions, which fit into a pattern similar to one of MGMT-positive hiPSCs. Blocking the initial stage of BER via MPG deficiency induces error-prone TLS through replication-blocking 3-meA and leads to the appearance of T>A substitutions. Remarkably, *MPG* knockout in the WT or *MSH2-/-* lines does not increase TMZ sensitivity, most probably because replication-blocking lesions are bypassed via TLS polymerases. However, interrupting BER at a later stage, e.g., in *XRCC1* knockout, leads to deletions, which likely indicate the presence of dsDNA breaks and cell death. Intriguingly, TLS through 3-meA results in different types of substitutions in BER-positive cells, which accumulate T>C transitions, and in BER-deficient cells, which, in addition to T>C transitions, accumulate a much larger number of T>A transversions. Different TLS polymerases may be involved in these lines. It is worth noting that T>C substitutions are unlikely to reflect the errors by BER DNA polymerase beta since their numbers and patterns do not change in *MSH2-/- MPG-/-* cells.

Our results suggest that targeting MPG or TLS polymerases will not lead to the re- sensitization of MMRd tumors to TMZ since BER and TLS act in parallel to ensure TMZ resistance. More promising targets appear to be XRCC1 and APE1 since initiation of BER at 3-meA and 7-meG leads to the conversion of these lesions into the abasic sites, nicks, and gaps, which become toxic if not processed promptly (Figure 8B). A recent report of a rare APE1-deficient glioblastoma which was effectively cured by the TMZ treatment, supports this view (79). It was suggested that XRCC1 releases PARP1 from ssDNA breaks, making them accessible for repair and preventing PARP1 hyperactivation. PARP inhibitors that efficiently trap PARP1 at the DNA breaks, such as olaparib and talazoparib, act similarly to XRCC1 deficiency and synergize with TMZ (58, 59). It appears that PARP-trapping compounds, as well as the potential future strategies/agents targeting XRCC1 and APE1, will be efficient primarily in preventing the acquisition of TMZ resistance via MMR deficiency but will not work on the primary MGMT-negative MMR-positive glioma since BER plays a minimal role in cell survival when the primary cytotoxic lesion is *O^6^*-meG. Targeting APE1 (80–82) appears to be a particularly promising strategy for potentiating the effect of TMZ in glioma treatment. The systematic analysis of isogenic series of repair-defective cells, combined with sensitivity measurements and signature analysis, holds great potential to uncover cancer cell vulnerabilities to genotoxic therapies.

## Methods

### Generation of cell lines

The following TK6 TSCER2(83)-derived cell lines were published previously: *EXO1-/-* (lab collection #24)(84), *FANCC-/-* (#27)(85), *FANCD2-/-* (#28)(86, 87), *LIG4-/-/-* (#34)(88), *MSH2-/-* (#52)(89), *MUS81-/-* (#54)(89), *PARP1-/-* (#64)(58), *PARP1-/- XRCC1-/-* (#65)(58), *POLB-/-* (#80)(90), *POLH-/-* (#84)(91), *POLI-/-* (#86)(35), *POLK-/-* (#88)(35), *RAD18-/-* (#93)(92), *XPA-/-* (#107)(91), *XRCC1-/-* (#97)(87), *XPA-/- XRCC1-/-* (#70)(93).

The triple knockout *POH-/- POLI-/- POLK-/-* (#181)(35) in TK6 TSCER2 was kindly provided by Prof. Kouji Hirota.

To generate knockout lines, we followed the procedure developed in Prof. Shunichi Takeda’s lab (89), which replaces one of the exons with antibiotic resistance markers, Neo, Hygro, Puro, or HisD. Two different resistance markers are used for the two alleles. Sequences of gRNAs and primers are listed in Supplementary Table 1. Overhangs used for cloning are shown in lowercase. Two complementary gRNA primers were annealed and cloned into the BbsI-digested pX330 vector (94). Two gRNAs targeting the same exon were used for each gene. An approximately 2 kb long genomic DNA fragment encompassing the exon of interest was PCR amplified with primers 2kb_F and 2kb_R and gel-purified. Left (LA) and right (RA) homology arms were then PCR amplified from the 2kb fragment with primer pairs LA_F and LA_R or RA_F and RA_R, respectively. The homology arms were cloned into ApaI/AflII-digested targeting vectors on either side of the resistance marker using GeneArt Seamless Cloning and Assembly Kit (Invitrogen). The targeting vectors, DT-A-pA/loxP/PGK-Neo (Hygro, Puro or HisD)-pA/loxP, were designed by the Laboratory for Animal Resources and Genetic Engineering, Center for Developmental Biology, RIKEN, Kobe (http://www.clst.riken.jp/arg). TK6 cells (∼6-8X10^6^) were electroporated in 100 microliter tips with 6 micrograms of gRNA/Cas9 and 4 micrograms of targeting plasmids using Neon Electroporation System (Invitrogen) following the Jurkat protocol. After 48 hours of recovery in an antibiotic-free medium, cells were seeded into media with corresponding antibiotics in 96-well plates. Antibiotic-resistant clones were expanded, and gene disruption was confirmed by PCR with the 2 kb_R primer, which anneals to the genomic DNA outside the right homology arm, and primer annealing to the resistance marker, NEO_F (AACCTGCGTGCAATCCATCTTGTTCAATGG), PURO_F (GTGAGGAAGAGTTCTTGCAGCTCGGTGA), HYGRO_F (ATCTTTGTAGAAACCATCGGCGCAGCTATT) or HIS_F (TTTATCAAATTTAGCGCTGTATTCACGCAG). The absence of the corresponding protein was confirmed by Western blot using the antibodies listed in the Supplementary Table 1.

To generate double knockouts *MSH2-/- ALKBH2-/-* (#172), *MSH2-/- ALKBH3-/-* (#171), *MSH2-/- MPG-/-* (#173), *MSH2-/- p53-/-* (#169), *MSH2-/- PARP1-/-* (#185), *MSH2-/- RAD18-/-* (#170), *MSH2-/- REV1-/-* (#192), *MSH2-/- REV7-/-* (#193), and *MSH2-/- XRCC1-/-* (#168), the *MSH2* gene was disrupted in the corresponding single knockout lines. To make *MSH2-/- APE1-/-* (#213), *MSH2-/- ATAD5-/-* (#161), and *MSH2-/- FANCD2-/-* (#155), *APE1, ATAD5*, and *FANCD2* genes, respectively, were knocked out in *MSH2-/-* parent. This change was implemented since *ATAD5-/-* and *FANCD2-/-* lines were slow-growing, and *APE1-/-* line was losing viability following electroporation.

To obtain *MSH2-/- XRCC1-/- MPG-/-* (#184) triple knockout, both alleles of the *MPG* gene were disrupted with Hygro marker in *MSH2-/- XRCC1-/-* line (#168), where *MSH2* gene had been knocked out using Neo and Puro and *XRCC1* gene using BSR and HisD markers.

To obtain *MSH2-/- XRCC1-/- PARP1-/-* (#197) triple knockout, Puro marker was looped out in the *XRCC1-/-PARP1-/-* line (#200) following transfection with the Cre-recombinase encoding plasmid. *MSH2* gene was then disrupted with Puro and Hygro markers.

To generate *RAD54L-/- RAD54B-/-* line (#210), we disrupted *RAD54L* gene in *RAD54B-/-* parent (a generous gift from Prof. Shunichi Takeda; #99).

HAP1 WT (C859, FACS-sorted for near-haploid status), *MGMT-* (20 bp deletion in exon 2; Product ID HZGHC000430c006) and *MSH2-* (2 bp deletion in exon 1; Product ID HZGHC024799c015) lines were obtained from Horizon. We confirmed that all the sequenced TMZ-treated subclones were mostly haploid by flow cytometry analysis.

### Clonogenic assays

TK6 cells were grown in RPMI1640 (Nacalai Tesque #30264-56) supplemented with 5% heat-inactivated donor horse serum (GIBCO #16050), 0.2 mg/ml Sodium Pyruvate and penicillin/streptomycin (Nacalai Tesque #09367-34). Temozolomide was purchased from Sigma-Aldrich (D4540) and dissolved in DMSO to make a 100 mM stock solution. Cells were diluted to 2X10^5^ cells/ml in 5 ml of medium in a 6-well plate and incubated for 24 hours, at which point TMZ was added, and cells were incubated for an additional 24 hours. DMSO was added to the control well instead of TMZ. Following incubation with the drug, cells were serially diluted to 1X10^4,^ and 1X10^3^ cells/ml and 200 microliters (∼2, 000 or 200 cells, respectively) were added to 5 ml of 1.5% methylcellulose (viscosity 1.500 cP, Sigma; M0387) pre-equilibrated in DMEM/HAM (Sigma; D8900) and 10% horse serum in a 6-well plate. Each experimental condition was replicated twice (technical replicates). Colonies were counted after 10-21 days of incubation, and the percentage of colonies in each TMZ concentration vs DMSO control was calculated. Clonogenic survival assays were repeated three times on different days (biological replicates) to determine the average surviving fraction and standard deviation.

HAP1 cells were grown in IMDM + GlutaMax^TM^ medium (ThermoFisher 31980-097) supplemented with 10% fetal bovine serum (Sigma TMS-013-BRK) and penicillin/streptomycin. For clonogenic survival assays, 1X10^3^ cells were seeded into 6 cm plates and incubated for 24 hours to allow them to adhere to the plate surface. Media was then changed to media with TMZ added to it. After 24 hr incubation with the drug, cells were washed with PBS and incubated in media without TMZ for six additional days. Colonies were fixed with 4% formaldehyde, stained with 1% methylene blue solution in 70% ethanol, and counted.

### Growing cell clones for whole-genome sequencing

TK6 cells were diluted to 1.5 cells/ml and seeded into 96-well plates at 200 microliters (∼0.3 cells) per well. After the cell number reached ∼2X10^5^ cells (17.5 doublings), the cells from one single clone were divided into 3 wells (50 microliters of cell suspension and 150 microliters of fresh media with or without TMZ). Following 24 hr incubation with the drug, cells were again diluted to 1.5 cells/ml and plated at 0.3 cells per well. Three subclones (∼2X10^5^ cells) per condition were harvested and sequenced.

Similar to TK6 cells, HAP1 clones were obtained by seeding 0.3 cells per well into 96 well plates. The selected clone was then expanded in 3 ml of media in 6-well plates until ∼ 4X10^6^ cells and seeded at ∼600, 000 cells/well in 3 ml of media in a 6-well plate for TMZ treatment on the following day. After 24 hr treatment with TMZ, cells were diluted to 0.3 cells/well and seeded into 96-well plates. Clones were then expanded in 3 ml of media in 6-well plates until the cell number reached ∼ 4X10^6^ and harvested. Half of the cells were used for sequencing and half for FACS analysis of the DNA content to confirm ploidy.

### Library construction and whole-genome sequencing

We performed whole-genome sequencing of wild-type HAP1 and TK6 cell lines and of 425 TMZ-treated and untreated samples of DNA repair mutants. DNA was extracted from cell pellets using the DNeasy Blood & Tissue Kit (Qiagen) according to the manufacturer’s instructions. The genomic DNA was treated with Frag enzyme to obtain DNA fragments between 100 and ∼1, 000 bp suitable for PE150 sequencing according to the manufacturer’s instructions (FS DNA library prep set, MGI, Shenzhen, China, cat No. 1, 000, 005, 256). The fragmented DNA was further selected to be between 300 and ∼500 bp by DNA clean beads (MGI, Shenzhen, China). The DNA fragments were blunt-ended, and dATP was added at the 3′ ends. The dTTP-tailed adapter sequence was ligated to both ends of the DNA fragments. The ligation product was then amplified for 7 cycles and subjected to the following single-strand circularization process. The PCR product was heat-denatured together with an oligonucleotide that was complementary to the adapter sequence, and the single-stranded circle ligated using DNA ligase. The remaining linear oligo was digested with the exonuclease, thus obtaining a single-strand circular DNA library. We sequenced the DNA library using DNBSEQ-T7 (RRID:SCR_017981) with a PE read length of 150 bp. Sequence depths were 100X for wild-type HAP1 and TK6 cell line controls and 30X for 425 TMZ-treated and untreated samples.

### WGS alignment

We used Burrows–Wheeler Aligner MEM (v.0.7.17) (95) tool with the human reference genome GRCh38/hg38 for WGS read alignment. The raw aligned reads were pre-processed using the GATK (v.4.1.9.0) (96) pipeline, including MarkDuplicates, BaseRecalibrator, and ApplyBQSR, to correct technical biases.

### Single nucleotide variant and short insertion and deletion detection

Single nucleotide variants (SNVs) and short insertion and deletions (indels) of subject samples were identified by Mutect2 (97) using untreated cell-line matched samples as controls The GATK Mutect2 workflow, including CalculateContamination, LearnReadOrientationModel, and FilterMutectCalls, was applied to filter out alignment artifacts and obtain somatic variants. Furthermore, additional filtering was performed to remove false positive calls. First, we selected SNVs and indels with a variant allele fraction greater than or equal to 0.2 with at least five variant-supporting reads. Second, we filtered out SNVs and indels with a read depth of less than 15 or greater than 150 at the variant site. Third, only the variants with at least one variant-supporting read in each read direction were further analyzed. Fourth, we used variants that were uniquely detected for individual samples.

### Structural variant identification

Delly2 (v.0.8.7) (98) was used to call structural variants (SVs). SVs were detected by comparing the subject samples to untreated cell-line matched controls. SVs that passed the default Delly2 quality filter (mapping quality ≧ 20 and paired-end support ≧ 5 reads for translocations or ≧ 3 reads for other SVs) were used for the subsequent analyses. We filtered out deletions and insertions less than 200bp long. We then selected only the unique variants that were found only once across all 425 samples.

### Extraction of mutational signatures

We utilized SigProfilerExtractor (v.1.1.4) (99) to extract de novo mutational signatures by a non-negative matrix factorization. The optimal number of mutational signatures was determined based on stability and cosine distance. Extracted de novo mutational signatures were reconstructed and decomposed to COSMIC mutational signatures.

### Microsatellite instability analysis

Microsatellite instability (MSI) indices were calculated using MANTIS (v.1.0.4) algorithm: cosine dissimilarity (cos), step-wise difference (dif), and Euclidean distance (euc) (53). The thresholds of instability recommended for the assessment of clinical samples are 0.07, 0.4, and 0.187 for cos, dif, and euc, respectively. Any value greater than or equal to the threshold is considered MSI unstable.

### Statistics and reproducibility

R software (v.4.0.3) (100) was employed for statistical analyses. The t-test used a compare_means function in the ggpubr package (100). A ci function in the bayestestR package (101) calculated 95 percent of the credible interval. The cosine correlation of mutational spectra was calculated by a cosine function in the lsa package (102). The number of variants of the samples used in the analysis is detailed in Supplementary Table 2.

## Data availability

The raw sequencing data are available for download from the Korean Nucleotide Archive (KoNA) under accession nos. KAP230727. All other data, including the number of variants of the samples, are available in the Supplementary Table 2.

## Code availability

The code used in this study is available.

## Acknowledgements

We are grateful to Kouji Hirota for sharing cell lines, to Orlando Schaerer for the many helpful discussions, and for reading and commenting on the manuscript, to Kyungjae Myung for his unwavering support, and to IBS-CGI members for a stimulating environment.

## Author contributions

DI and AG conceptualization, supervision, data interpretation, and writing. SL supervision of all bioinformatics work and funding acquisition. TH all bioinformatics work, data interpretation, and generation of figures. JW software and hardware setup. ST provision of cell lines and sharing experimental procedures. LKS data interpretation. DI, LKS, RK, FNA, DMS, HP, BC, and LM experimental work.

## References

1. Alexandrov, L.B., Nik-Zainal, S., Wedge, D.C., Aparicio, S.A.J.R., Behjati, S., Biankin, A.V., Bignell, G.R., Bolli, N., Borg, A., Børresen-Dale, A.-L., et al. (2013) Signatures of mutational processes in human cancer. Nature, 500, 415–421.

2. Kucab, J.E., Zou, X., Morganella, S., Joel, M., Nanda, A.S., Nagy, E., Gomez, C., Degasperi, A., Harris, R., Jackson, S.P., et al. (2019) A Compendium of Mutational Signatures of Environmental Agents. Cell, 177, 821–836.e16.

3. Zou, X., Koh, G.C.C., Nanda, A.S., Degasperi, A., Urgo, K., Roumeliotis, T.I., Agu, C.A., Badja, C., Momen, S., Young, J., et al. (2021) A systematic CRISPR screen defines mutational mechanisms underpinning signatures caused by replication errors and endogenous DNA damage. Nat Cancer, 2, 643–657.

4. Zou, X., Owusu, M., Harris, R., Jackson, S.P., Loizou, J.I. and Nik-Zainal, S. (2018) Validating the concept of mutational signatures with isogenic cell models. Nat. Commun., 9, 1744.

5. Volkova, N.V., Meier, B., González-Huici, V., Bertolini, S., Gonzalez, S., Vöhringer, H., Abascal, F., Martincorena, I., Campbell, P.J., Gartner, A., et al. (2020) Mutational signatures are jointly shaped by DNA damage and repair. Nat. Commun., 11, 2169.

6. Cagan, A., Baez-Ortega, A., Brzozowska, N., Abascal, F., Coorens, T.H.H., Sanders, M.A., Lawson, A.R.J., Harvey, L.M.R., Bhosle, S., Jones, D., et al. (2022) Somatic mutation rates scale with lifespan across mammals. Nature, 604, 517–524.

7. Alexandrov, L.B., Jones, P.H., Wedge, D.C., Sale, J.E., Campbell, P.J., Nik-Zainal, S. and Stratton, M.R. (2015) Clock-like mutational processes in human somatic cells. Nat. Genet., 47, 1402–1407.

8. Kaina, B. and Christmann, M. (2019) DNA repair in personalized brain cancer therapy with temozolomide and nitrosoureas. DNA Repair, 78, 128–141.

9. Fu, D., Calvo, J.A. and Samson, L.D. (2012) Balancing repair and tolerance of DNA damage caused by alkylating agents. Nat. Rev. Cancer, 12, 104–120.

10. Denny, B.J., Wheelhouse, R.T., Stevens, M.F., Tsang, L.L. and Slack, J.A. (1994) NMR and molecular modeling investigation of the mechanism of activation of the antitumor drug temozolomide and its interaction with DNA. Biochemistry, 33, 9045–9051.

11. Christmann, M., Verbeek, B., Roos, W.P. and Kaina, B. (2011) O6-Methylguanine-DNA methyltransferase (MGMT) in normal tissues and tumors: Enzyme activity, promoter methylation and immunohistochemistry. Biochimica et Biophysica Acta (BBA) - Reviews on Cancer, 1816, 179–190.

12. Hegi, M.E., Diserens, A.-C., Gorlia, T., Hamou, M.-F., de Tribolet, N., Weller, M., Kros, J.M., Hainfellner, J.A., Mason, W., Mariani, L., et al. (2005) MGMT gene silencing and benefit from temozolomide in glioblastoma. N. Engl. J. Med., 352, 997–1003.

13. Karran, P. and Bignami, M. (1994) DNA damage tolerance, mismatch repair and genome instability. Bioessays, 16, 833–839.

14. Stojic, L., Mojas, N., Cejka, P., Di Pietro, M., Ferrari, S., Marra, G. and Jiricny, J. (2004) Mismatch repair-dependent G2 checkpoint induced by low doses of SN1 type methylating agents requires the ATR kinase. Genes Dev., 18, 1331–1344.

15. Plant, J.E. and Roberts, J.J. (1971) A novel mechanism for the inhibition of DNA synthesis following methylation: the effect of N-methyl-N-nitrosourea on HeLa cells. Chem. Biol. Interact., 3, 337–342.

16. Yoshioka, K.-I., Yoshioka, Y. and Hsieh, P. (2006) ATR kinase activation mediated by MutSalpha and MutLalpha in response to cytotoxic O6-methylguanine adducts. Mol. Cell, 22, 501–510.

17. Lin, B., Gupta, D. and Heinen, C.D. (2014) Human pluripotent stem cells have a novel mismatch repair-dependent damage response. J. Biol. Chem., 289, 24314–24324.

18. Gupta, D., Lin, B., Cowan, A. and Heinen, C.D. (2018) ATR-Chk1 activation mitigates replication stress caused by mismatch repair-dependent processing of DNA damage. Proc. Natl. Acad. Sci. U. S. A., 115, 1523–1528.

19. Fuchs, R.P., Isogawa, A., Paulo, J.A., Onizuka, K., Takahashi, T., Amunugama, R., Duxin, J.P. and Fujii, S. (2021) Crosstalk between repair pathways elicits double-strand breaks in alkylated DNA and implications for the action of temozolomide. Elife, 10.

20. Rinne, M.L., He, Y., Pachkowski, B.F., Nakamura, J. and Kelley, M.R. (2005) N-methylpurine DNA glycosylase overexpression increases alkylation sensitivity by rapidly removing non-toxic 7-methylguanine adducts. Nucleic Acids Res., 33, 2859–2867.

21. Wilson, D.M., Iii (2016) Base Excision Repair Pathway, The: Molecular Mechanisms And Role In Disease Development And Therapeutic Design World Scientific.

22. Plosky, B., Samson, L., Engelward, B.P., Gold, B., Schlaen, B., Millas, T., Magnotti, M., Schor, J. and Scicchitano, D.A. (2002) Base excision repair and nucleotide excision repair contribute to the removal of N-methylpurines from active genes. DNA Repair, 1, 683–696.

23. Monti, P., Iannone, R., Campomenosi, P., Ciribilli, Y., Varadarajan, S., Shah, D., Menichini, P., Gold, B. and Fronza, G. (2004) Nucleotide excision repair defect influences lethality and mutagenicity induced by Me-lex, a sequence-selective N3-adenine methylating agent in the absence of base excision repair. Biochemistry, 43, 5592–5599.

24. Monti, P., Ciribilli, Y., Russo, D., Bisio, A., Perfumo, C., Andreotti, V., Menichini, P., Inga, A., Huang, X., Gold, B., et al. (2008) Rev1 and Polzeta influence toxicity and mutagenicity of Me-lex, a sequence selective N3-adenine methylating agent. DNA Repair, 7, 431–438.

25. Póti, Á., Szikriszt, B., Gervai, J.Z., Chen, D. and Szüts, D. (2022) Characterisation of the spectrum and genetic dependence of collateral mutations induced by translesion DNA synthesis. PLoS Genet., 18, e1010051.

26. Szikriszt, B., Póti, Á., Németh, E., Kanu, N., Swanton, C. and Szüts, D. (2021) A comparative analysis of the mutagenicity of platinum-containing chemotherapeutic agents reveals direct and indirect mutagenic mechanisms. Mutagenesis, 36, 75–86.

27. Kuijk, E., Jager, M., van der Roest, B., Locati, M.D., Van Hoeck, A., Korzelius, J., Janssen, R., Besselink, N., Boymans, S., van Boxtel, R., et al. (2020) The mutational impact of culturing human pluripotent and adult stem cells. Nat. Commun., 11, 2493.

28. Alexandrov, L.B., Kim, J., Haradhvala, N.J., Huang, M.N., Tian Ng, A.W., Wu, Y., Boot, A., Covington, K.R., Gordenin, D.A., Bergstrom, E.N., et al. (2020) The repertoire of mutational signatures in human cancer. Nature, 578, 94–101.

29. Abascal, F., Harvey, L.M.R., Mitchell, E., Lawson, A.R.J., Lensing, S.V., Ellis, P., Russell, A.J.C., Alcantara, R.E., Baez-Ortega, A., Wang, Y., et al. (2021) Somatic mutation landscapes at single-molecule resolution. Nature, 593, 405–410.

30. Koh, G., Degasperi, A., Zou, X., Momen, S. and Nik-Zainal, S. (2021) Mutational signatures: emerging concepts, caveats and clinical applications. Nat. Rev. Cancer, 21, 619–637.

31. Zámborszky, J., Szikriszt, B., Gervai, J.Z., Pipek, O., Póti, Á., Krzystanek, M., Ribli, D., Szalai-Gindl, J.M., Csabai, I., Szallasi, Z., et al. (2017) Loss of BRCA1 or BRCA2 markedly increases the rate of base substitution mutagenesis and has distinct effects on genomic deletions. Oncogene, 36, 5085–5086.

32. Makarova, A.V. and Burgers, P.M. (2015) Eukaryotic DNA polymerase ζ. DNA Repair, 29, 47–55.

33. van Bostelen, I., van Schendel, R., Romeijn, R. and Tijsterman, M. (2020) Translesion synthesis polymerases are dispensable for C. elegans reproduction but suppress genome scarring by polymerase theta-mediated end joining. PLoS Genet., 16, e1008759.

34. Meier, B., Volkova, N.V., Hong, Y., Bertolini, S., González-Huici, V., Petrova, T., Boulton, S., Campbell, P.J., Gerstung, M. and Gartner, A. (2021) Protection of the C. elegans germ cell genome depends on diverse DNA repair pathways during normal proliferation. PLoS One, 16, e0250291.

35. Inomata, Y., Abe, T., Tsuda, M., Takeda, S. and Hirota, K. (2021) Division of labor of Y-family polymerases in translesion-DNA synthesis for distinct types of DNA damage. PLoS One, 16, e0252587.

36. Szüts, D., Marcus, A.P., Himoto, M., Iwai, S. and Sale, J.E. (2008) REV1 restrains DNA polymerase zeta to ensure frame fidelity during translesion synthesis of UV photoproducts in vivo. Nucleic Acids Res., 36, 6767–6780.

37. Wittschieben, J.P., Reshmi, S.C., Gollin, S.M. and Wood, R.D. (2006) Loss of DNA polymerase zeta causes chromosomal instability in mammalian cells. Cancer Res., 66, 134–142.

38. Ivanov, D., Hwang, T., Sitko, L.K., Lee, S. and Gartner, A. (2023) Experimental systems for the analysis of mutational signatures: no ‘one-size-fits-all’ solution. Biochem. Soc. Trans., 51, 1307–1317.

39. Korona, D.A., Lecompte, K.G. and Pursell, Z.F. (2011) The high fidelity and unique error signature of human DNA polymerase epsilon. Nucleic Acids Res., 39, 1763–1773.

40. Schmitt, M.W., Matsumoto, Y. and Loeb, L.A. (2009) High fidelity and lesion bypass capability of human DNA polymerase delta. Biochimie, 91, 1163–1172.

41. Bartsch, C., Bartsch, H., Blask, D.E., Cardinali, D.P., Hrushesky, W.J.M. and Mecke, D. (2012) The Pineal Gland and Cancer: Neuroimmunoendocrine Mechanisms in Malignancy Springer Science & Business Media.

42. Lancey, C., Tehseen, M., Raducanu, V.-S., Rashid, F., Merino, N., Ragan, T.J., Savva, C.G., Zaher, M.S., Shirbini, A., Blanco, F.J., et al. (2020) Structure of the processive human Pol δ holoenzyme. Nat. Commun., 11, 1109.

43. Langston, L.D., Zhang, D., Yurieva, O., Georgescu, R.E., Finkelstein, J., Yao, N.Y., Indiani, C. and O’Donnell, M.E. (2014) CMG helicase and DNA polymerase ε form a functional 15-subunit holoenzyme for eukaryotic leading-strand DNA replication. Proc. Natl. Acad. Sci. U. S. A., 111, 15390–15395.

44. Zhou, J.C., Janska, A., Goswami, P., Renault, L., Abid Ali, F., Kotecha, A., Diffley, J.F.X. and Costa, A. (2017) CMG-Pol epsilon dynamics suggests a mechanism for the establishment of leading-strand synthesis in the eukaryotic replisome. Proc. Natl. Acad. Sci. U. S. A., 114, 4141–4146.

45. Dmowski, M., Makiela-Dzbenska, K., Sharma, S., Chabes, A. and Fijalkowska, I.J. (2023) Impairment of the non-catalytic subunit Dpb2 of DNA Pol L results in increased involvement of Pol δ on the leading strand. DNA Repair, 129, 103541.

46. Grabowska, E., Wronska, U., Denkiewicz, M., Jaszczur, M., Respondek, A., Alabrudzinska, M., Suski, C., Makiela-Dzbenska, K., Jonczyk, P. and Fijalkowska, I.J. (2014) Proper functioning of the GINS complex is important for the fidelity of DNA replication in yeast. Mol. Microbiol., 92, 659–680.

47. Lózsa, R., Németh, E., Gervai, J.Z., Márkus, B.G., Kollarics, S., Gyüre, Z., Tóth, J., Simon, F. and Szüts, D. (2023) DNA mismatch repair protects the genome from oxygen-induced replicative mutagenesis. Nucleic Acids Res., 51, 11040–11055.

48. Thibodeau, S.N., Bren, G. and Schaid, D. (1993) Microsatellite instability in cancer of the proximal colon. Science, 260, 816–819.

49. Touat, M., Li, Y.Y., Boynton, A.N., Spurr, L.F., Iorgulescu, J.B., Bohrson, C.L., Cortes-Ciriano, I., Birzu, C., Geduldig, J.E., Pelton, K., et al. (2020) Mechanisms and therapeutic implications of hypermutation in gliomas. Nature, 580, 517–523.

50. Hayashida, G., Shioi, S., Hidaka, K., Fujikane, R., Hidaka, M., Tsurimoto, T., Tsuzuki, T., Oda, S. and Nakatsu, Y. (2019) Differential genomic destabilisation in human cells with pathogenic MSH2 mutations introduced by genome editing. Exp. Cell Res., 377, 24–35.

51. Boland, C.R., Thibodeau, S.N., Hamilton, S.R., Sidransky, D., Eshleman, J.R., Burt, R.W., Meltzer, S.J., Rodriguez-Bigas, M.A., Fodde, R., Ranzani, G.N., et al. (1998) A National Cancer Institute Workshop on Microsatellite Instability for cancer detection and familial predisposition: development of international criteria for the determination of microsatellite instability in colorectal cancer. Cancer Res., 58, 5248–5257.

52. Hempelmann, J.A., Lockwood, C.M., Konnick, E.Q., Schweizer, M.T., Antonarakis, E.S., Lotan, T.L., Montgomery, B., Nelson, P.S., Klemfuss, N., Salipante, S.J., et al. (2018) Microsatellite instability in prostate cancer by PCR or next-generation sequencing. J Immunother Cancer, 6, 29.

53. Kautto, E.A., Bonneville, R., Miya, J., Yu, L., Krook, M.A., Reeser, J.W. and Roychowdhury, S. (2017) Performance evaluation for rapid detection of pan-cancer microsatellite instability with MANTIS. Oncotarget, 8, 7452–7463.

54. Webster, A.L.H., Sanders, M.A., Patel, K., Dietrich, R., Noonan, R.J., Lach, F.P., White, R.R., Goldfarb, A., Hadi, K., Edwards, M.M., et al. (2022) Genomic signature of Fanconi anaemia DNA repair pathway deficiency in cancer. Nature, 612, 495–502.

55. Mórocz, M., Qorri, E., Pekker, E., Tick, G. and Haracska, L. (2024) Exploring RAD18- dependent replication of damaged DNA and discontinuities: A collection of advanced tools. J. Biotechnol., 380, 1–19.

56. MacLeod, G., Bozek, D.A., Rajakulendran, N., Monteiro, V., Ahmadi, M., Steinhart, Z., Kushida, M.M., Yu, H., Coutinho, F.J., Cavalli, F.M.G., et al. (2019) Genome-Wide CRISPR-Cas9 Screens Expose Genetic Vulnerabilities and Mechanisms of Temozolomide Sensitivity in Glioblastoma Stem Cells. Cell Rep., 27, 971–986.e9.

57. Caldecott, K.W. (2019) XRCC1 protein; Form and function. DNA Repair, 81, 102664.

58. Demin, A.A., Hirota, K., Tsuda, M., Adamowicz, M., Hailstone, R., Brazina, J., Gittens, W., Kalasova, I., Shao, Z., Zha, S., et al. (2021) XRCC1 prevents toxic PARP1 trapping during DNA base excision repair. Mol. Cell, 81, 3018–3030.e5.

59. Hirota, K., Ooka, M., Shimizu, N., Yamada, K., Tsuda, M., Ibrahim, M.A., Yamada, S., Sasanuma, H., Masutani, M. and Takeda, S. (2022) XRCC1 counteracts poly(ADP ribose)polymerase (PARP) poisons, olaparib and talazoparib, and a clinical alkylating agent, temozolomide, by promoting the removal of trapped PARP1 from broken DNA. Genes Cells, 27, 331–344.

60. Bryant, H.E., Petermann, E., Schultz, N., Jemth, A.-S., Loseva, O., Issaeva, N., Johansson, F., Fernandez, S., McGlynn, P. and Helleday, T. (2009) PARP is activated at stalled forks to mediate Mre11-dependent replication restart and recombination. EMBO J., 28, 2601–2615.

61. Hanzlikova, H., Kalasova, I., Demin, A.A., Pennicott, L.E., Cihlarova, Z. and Caldecott, K.W. (2018) The Importance of Poly(ADP-Ribose) Polymerase as a Sensor of Unligated Okazaki Fragments during DNA Replication. Mol. Cell, 71, 319–331.e3.

62. Schlacher, K., Wu, H. and Jasin, M. (2012) A distinct replication fork protection pathway connects Fanconi anemia tumor suppressors to RAD51-BRCA1/2. Cancer Cell, 22, 106–116.

63. Olivieri, M., Cho, T., Álvarez-Quilón, A., Li, K., Schellenberg, M.J., Zimmermann, M., Hustedt, N., Rossi, S.E., Adam, S., Melo, H., et al. (2020) A Genetic Map of the Response to DNA Damage in Human Cells. Cell, 182, 481–496.e21.

64. Peng, M., Xie, J., Ucher, A., Stavnezer, J. and Cantor, S.B. (2014) Crosstalk between BRCA-Fanconi anemia and mismatch repair pathways prevents MSH2-dependent aberrant DNA damage responses. EMBO J., 33, 1698–1712.

65. Paul Solomon Devakumar, L.J., Gaubitz, C., Lundblad, V., Kelch, B.A. and Kubota, T. (2019) Effective mismatch repair depends on timely control of PCNA retention on DNA by the Elg1 complex. Nucleic Acids Res., 47, 6826–6841.

66. Lee, K.-Y., Fu, H., Aladjem, M.I. and Myung, K. (2013) ATAD5 regulates the lifespan of DNA replication factories by modulating PCNA level on the chromatin. J. Cell Biol., 200, 31–44.

67. Willis, N.A., Frock, R.L., Menghi, F., Duffey, E.E., Panday, A., Camacho, V., Hasty, E.P., Liu, E.T., Alt, F.W. and Scully, R. (2017) Mechanism of tandem duplication formation in BRCA1-mutant cells. Nature, 551, 590–595.

68. Menghi, F., Inaki, K., Woo, X., Kumar, P.A., Grzeda, K.R., Malhotra, A., Yadav, V., Kim, H., Marquez, E.J., Ucar, D., et al. (2016) The tandem duplicator phenotype as a distinct genomic configuration in cancer. Proc. Natl. Acad. Sci. U. S. A., 113, E2373–82.

69. Nik-Zainal, S., Davies, H., Staaf, J., Ramakrishna, M., Glodzik, D., Zou, X., Martincorena, I., Alexandrov, L.B., Martin, S., Wedge, D.C., et al. (2016) Landscape of somatic mutations in 560 breast cancer whole-genome sequences. Nature, 534, 47–54.

70. Park, S.H., Kim, N., Kang, N., Ryu, E., Lee, E.A., Ra, J.S., Gartner, A., Kang, S., Myung, K. and Lee, K.-Y. (2023) Short-range end resection requires ATAD5-mediated PCNA unloading for faithful homologous recombination. Nucleic Acids Res., 51, 10519–10535.

71. Johannessen, T.-C.A., Prestegarden, L., Grudic, A., Hegi, M.E., Tysnes, B.B. and Bjerkvig, R. (2013) The DNA repair protein ALKBH2 mediates temozolomide resistance in human glioblastoma cells. Neuro. Oncol., 15, 269–278.

72. Dango, S., Mosammaparast, N., Sowa, M.E., Xiong, L.-J., Wu, F., Park, K., Rubin, M., Gygi, S., Harper, J.W. and Shi, Y. (2011) DNA unwinding by ASCC3 helicase is coupled to ALKBH3-dependent DNA alkylation repair and cancer cell proliferation. Mol. Cell, 44, 373–384.

73. Calvo, J.A., Meira, L.B., Lee, C.-Y.I., Moroski-Erkul, C.A., Abolhassani, N., Taghizadeh, K., Eichinger, L.W., Muthupalani, S., Nordstrand, L.M., Klungland, A., et al. (2012) DNA repair is indispensable for survival after acute inflammation. J. Clin. Invest., 122, 2680–2689.

74. Haghani, A., Li, C.Z., Robeck, T.R., Zhang, J., Lu, A.T., Ablaeva, J., Acosta-Rodríguez, V.A., Adams, D.M., Alagaili, A.N., Almunia, J., et al. (2023) DNA methylation networks underlying mammalian traits. Science, 381, eabq5693.

75. Lu, A.T., Fei, Z., Haghani, A., Robeck, T.R., Zoller, J.A., Li, C.Z., Lowe, R., Yan, Q., Zhang, J., Vu, H., et al. (2023) Universal DNA methylation age across mammalian tissues. Nat Aging, 3, 1144–1166.

76. Gyüre, Z., Póti, Á., Németh, E., Szikriszt, B., Lózsa, R., Krawczyk, M., Richardson, A.L. and Szüts, D. (2023) Spontaneous mutagenesis in human cells is controlled by REV1- Polymerase ζ and PRIMPOL. Cell Rep., 42, 112887.

77. Tomasetti, C., Marchionni, L., Nowak, M.A., Parmigiani, G. and Vogelstein, B. (2015) Only three driver gene mutations are required for the development of lung and colorectal cancers. Proc. Natl. Acad. Sci. U. S. A., 112, 118–123.

78. Hartley, J.A., Mattes, W.B., Vaughan, K. and Gibson, N.W. (1988) DNA sequence specificity of guanine N7-alkylations for a series of structurally related triazenes. Carcinogenesis, 9, 669–674.

79. Wheeler, D.A., Takebe, N., Hinoue, T., Hoadley, K.A., Cardenas, M.F., Hamilton, A.M., Laird, P.W., Wang, L., Johnson, A., Dewal, N., et al. (2021) Molecular Features of Cancers Exhibiting Exceptional Responses to Treatment. Cancer Cell, 39, 38–53.e7.

80. Malfatti, M.C., Bellina, A., Antoniali, G. and Tell, G. (2023) Revisiting Two Decades of Research Focused on Targeting APE1 for Cancer Therapy: The Pros and Cons. Cells, 12.

81. Wilson, D.M., 3rd, Deacon, A.M., Duncton, M.A.J., Pellicena, P., Georgiadis, M.M., Yeh, A.P., Arvai, A.S., Moiani, D., Tainer, J.A. and Das, D. (2021) Fragment- and structure-based drug discovery for developing therapeutic agents targeting the DNA Damage Response. Prog. Biophys. Mol. Biol., 163, 130–142.

82. Caston, R.A., Gampala, S., Armstrong, L., Messmann, R.A., Fishel, M.L. and Kelley, M.R. (2021) The multifunctional APE1 DNA repair-redox signaling protein as a drug target in human disease. Drug Discov. Today, 26, 218–228.

83. Honma, M., Izumi, M., Sakuraba, M., Tadokoro, S., Sakamoto, H., Wang, W., Yatagai, F. and Hayashi, M. (2003) Deletion, rearrangement, and gene conversion; genetic consequences of chromosomal doubleLstrand breaks in human cells. Environ. Mol. Mutagen., 42, 288–298.

84. Kratz, K., Artola-Borán, M., Kobayashi-Era, S., Koh, G., Oliveira, G., Kobayashi, S., Oliveira, A., Zou, X., Richter, J., Tsuda, M., et al. (2021) FANCD2-Associated Nuclease 1 Partially Compensates for the Lack of Exonuclease 1 in Mismatch Repair. Mol. Cell. Biol., 41, e0030321.

85. Xu, X., Xu, Y., Guo, R., Xu, R., Fu, C., Xing, M., Sasanuma, H., Li, Q., Takata, M., Takeda, S., et al. (2021) Fanconi anemia proteins participate in a break-induced-replication-like pathway to counter replication stress. Nat. Struct. Mol. Biol., 28, 487–500.

86. Hashimoto, K., Sharma, V., Sasanuma, H., Tian, X., Takata, M., Takeda, S., Swenberg, J.A. and Nakamura, J. (2016) Poor recognition of O6-isopropyl dG by MGMT triggers double strand break-mediated cell death and micronucleus induction in FANC-deficient cells. Oncotarget, 7, 59795–59808.

87. Saha, L.K., Kim, S., Kang, H., Akter, S., Choi, K., Sakuma, T., Yamamoto, T., Sasanuma, H., Hirota, K., Nakamura, J., et al. (2018) Differential micronucleus frequency in isogenic human cells deficient in DNA repair pathways is a valuable indicator for evaluating genotoxic agents and their genotoxic mechanisms. Environ. Mol. Mutagen., 59, 529– 538.

88. Keka, I.S., Mohiuddin, Maede, Y., Rahman, M.M., Sakuma, T., Honma, M., Yamamoto, T., Takeda, S. and Sasanuma, H. (2015) Smarcal1 promotes double-strand-break repair by nonhomologous end-joining. Nucleic Acids Res., 43, 6359–6372.

89. Rahman, M.M., Mohiuddin, M., Shamima Keka, I., Yamada, K., Tsuda, M., Sasanuma, H., Andreani, J., Guerois, R., Borde, V., Charbonnier, J.-B., et al. (2020) Genetic evidence for the involvement of mismatch repair proteins, PMS2 and MLH3, in a late step of homologous recombination. J. Biol. Chem., 295, 17460–17475.

90. Saha, L.K., Wakasugi, M., Akter, S., Prasad, R., Wilson, S.H., Shimizu, N., Sasanuma, H., Huang, S.-Y.N., Agama, K., Pommier, Y., et al. (2020) Topoisomerase I-driven repair of UV-induced damage in NER-deficient cells. Proc. Natl. Acad. Sci. U. S. A., 117, 14412– 14420.

91. Mohiuddin, M., Evans, T.J., Rahman, M.M., Keka, I.S., Tsuda, M., Sasanuma, H. and Takeda, S. (2018) SUMOylation of PCNA by PIAS1 and PIAS4 promotes template switch in the chicken and human B cell lines. Proc. Natl. Acad. Sci. U. S. A., 115, 12793–12798.

92. Tsuda, M., Terada, K., Ooka, M., Kobayashi, K., Sasanuma, H., Fujisawa, R., Tsurimoto, T., Yamamoto, J., Iwai, S., Kadoda, K., et al. (2017) The dominant role of proofreading exonuclease activity of replicative polymerase ε in cellular tolerance to cytarabine (Ara- C). Oncotarget, 8, 33457–33474.

93. Ibrahim, M.A., Yasui, M., Saha, L.K., Sasanuma, H., Honma, M. and Takeda, S. (2020) Enhancing the sensitivity of the thymidine kinase assay by using DNA repair-deficient human TK6 cells. Environ. Mol. Mutagen., 61, 602–610.

94. Cong, L., Ran, F.A., Cox, D., Lin, S., Barretto, R., Habib, N., Hsu, P.D., Wu, X., Jiang, W., Marraffini, L.A., et al. (2013) Multiplex genome engineering using CRISPR/Cas systems. Science, 339, 819–823.

95. Li, H. (2013) Aligning sequence reads, clone sequences and assembly contigs with BWA-MEM. arXiv [q-bio.GN].

96. McKenna, A., Hanna, M., Banks, E., Sivachenko, A., Cibulskis, K., Kernytsky, A., Garimella, K., Altshuler, D., Gabriel, S., Daly, M., et al. (2010) The Genome Analysis Toolkit: a MapReduce framework for analyzing next-generation DNA sequencing data. Genome Res., 20, 1297–1303.

97. Benjamin, D., Sato, T., Cibulskis, K., Getz, G., Stewart, C. and Lichtenstein, L. (2019) Calling Somatic SNVs and Indels with Mutect2. bioRxiv, 10.1101/861054.

98. Rausch, T., Zichner, T., Schlattl, A., Stütz, A.M., Benes, V. and Korbel, J.O. (2012) DELLY: structural variant discovery by integrated paired-end and split-read analysis. Bioinformatics, 28, i333–i339.

99. Islam, S.M.A., Díaz-Gay, M., Wu, Y., Barnes, M., Vangara, R., Bergstrom, E.N., He, Y., Vella, M., Wang, J., Teague, J.W., et al. (2022) Uncovering novel mutational signatures by de novo extraction with SigProfilerExtractor. Cell Genom, 2, None.

100. Team, R.D.C. (2010) R: A language and environment for statistical computing. (No Title).

