## Supplementary Figures for "Comprehensive Whole Genome Sequencing Reveals Origins of Mutational Signatures Associated with Aging and Temozolomide Chemotherapy"

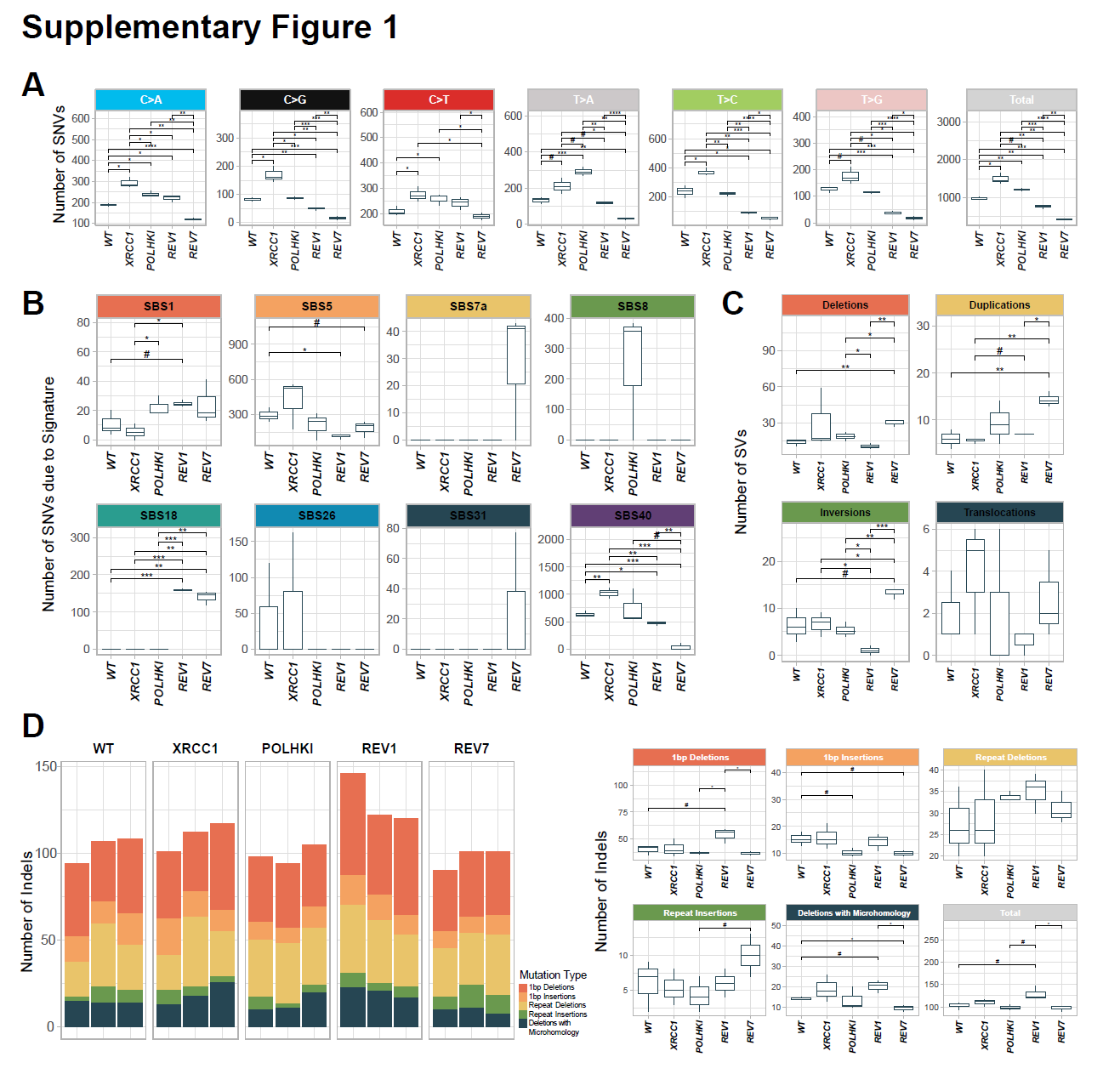
**Supplementary Figure S1.** Box plots of spontaneous mutations. Box plots indicate the statistical significance of differences in mutation numbers among the cell lines in Figure 2. Panels **A**, **B**, and **C** correspond to panels **A**, **C**, and **D** of Figure 2, respectively. **(D)** Numbers of indels accumulated in different cell lines after ~180 cell generations (10 passages). Significant differences are indicated by # and * (# p value ≤0.1, * p value ≤ 0.05, ** p value ≤ 0.01, *** p value ≤ 0.001, t-test).

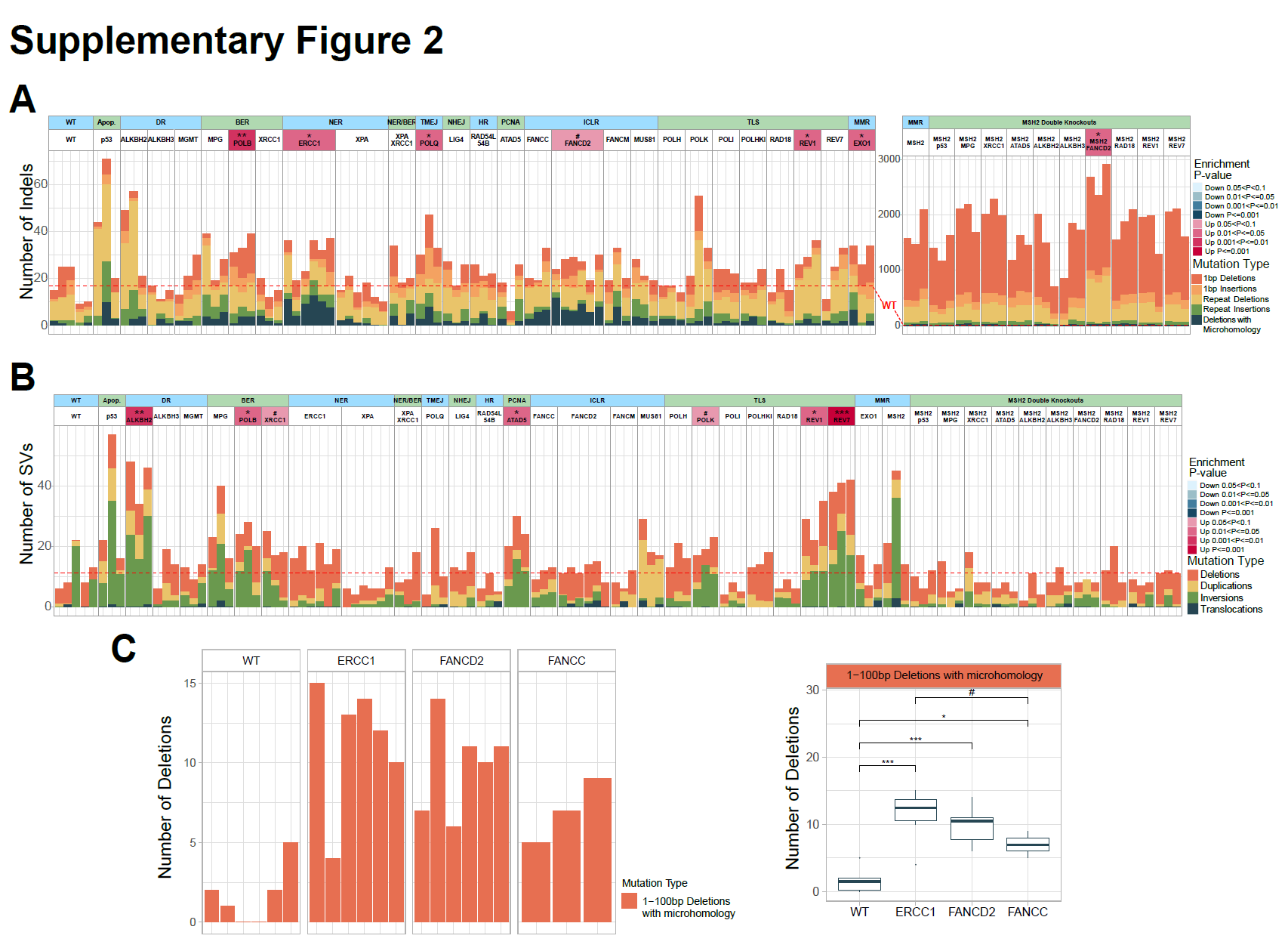
**Supplementary Figure S2.** Spontaneous mutagenesis in the collection of TK6 DNA repair mutants. Numbers of indels **(A)** and SVs **(B)** accumulated after 18 cell generations in individual subclones. The dashed line corresponds to an average of WT. The statistical tests on the left panel are compared to the WT lines and the right panel is compared to the *MSH2-/-* lines. **(C)** Deletions with microhomology in *ERCC1-/-*, *FANCD2-/-*, and *FANCC-/-* lines. Numbers of 1-100 bp deletions flanked by microhomologies. Significant differences are indicated by # and * (# p value ≤0.1, * p value ≤ 0.05, ** p value ≤ 0.01, *** p value ≤ 0.001, t-test).

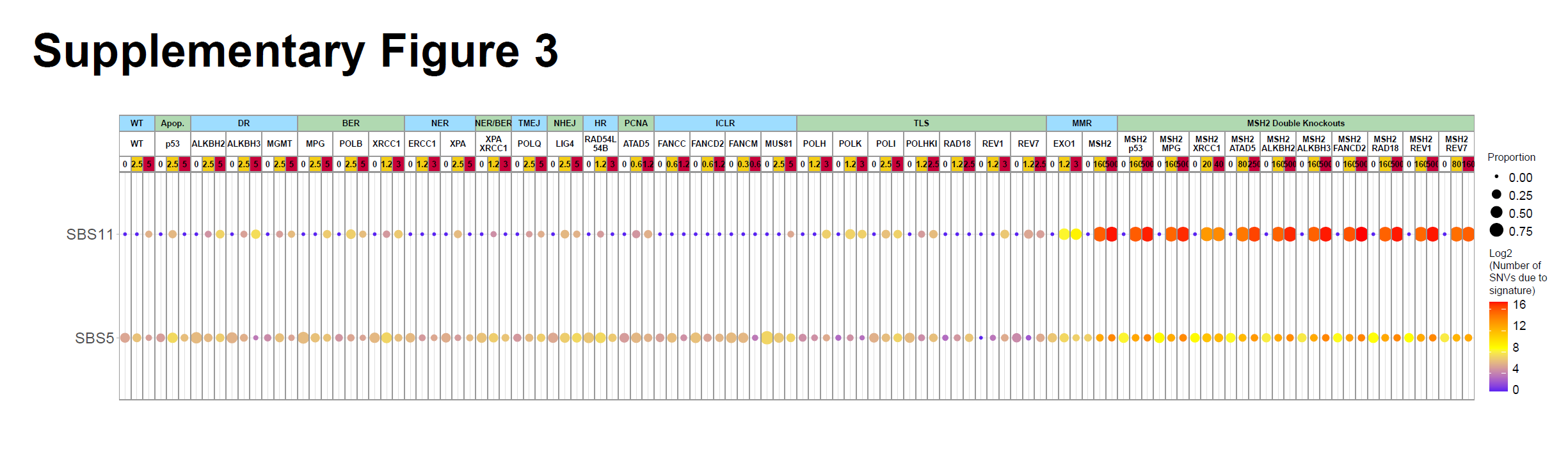
**Supplementary Figure S3.** Signatures SBS5 and SBS11 induced by TMZ. Prominence of signatures SBS5 and SBS11 in the TMZ-induced mutational profiles. The size of the dot represents the proportion of SNVs, and the color represents the log2 of the number of SNVs due to signature SBS5 or SBS11.

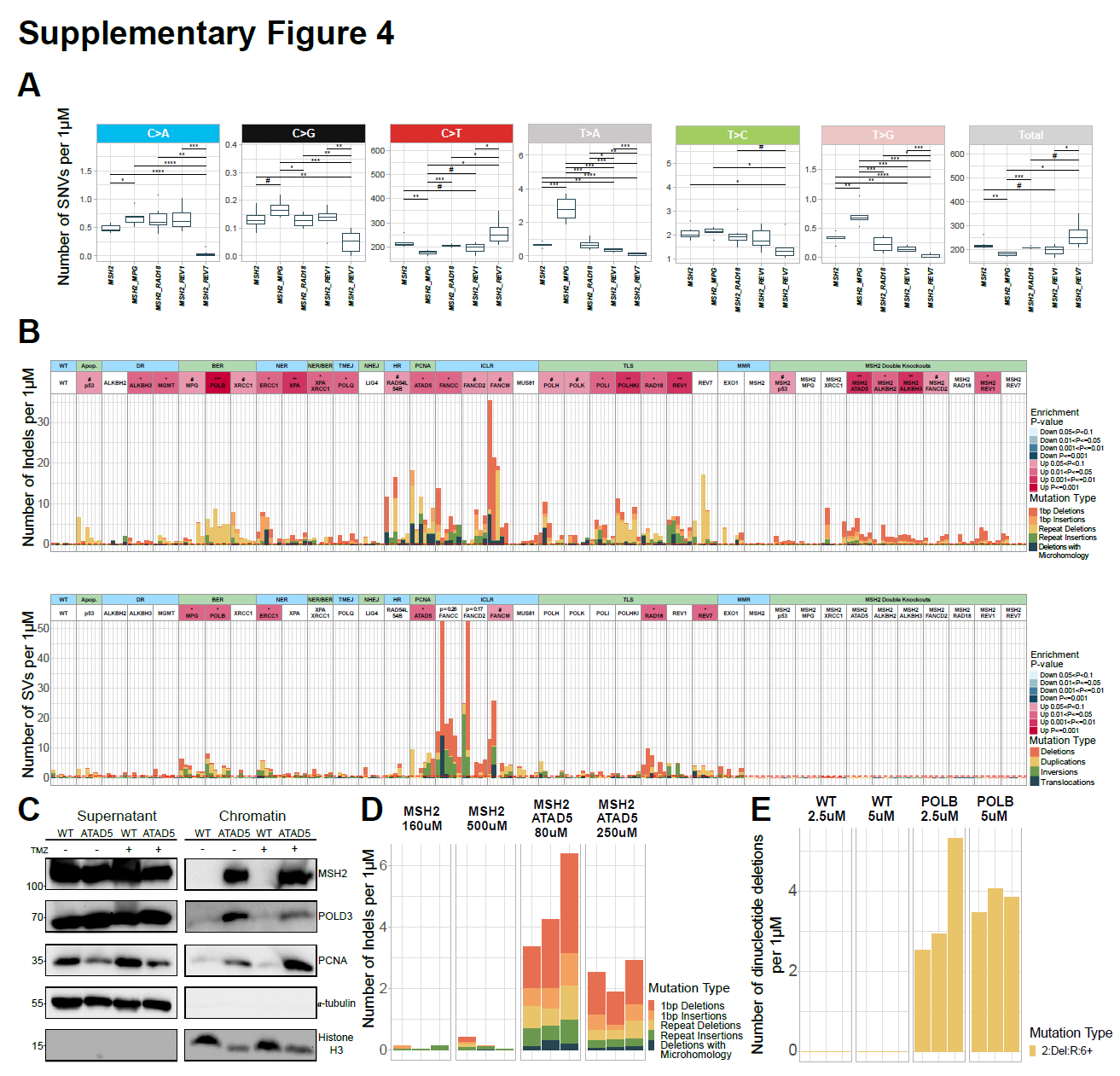
**Supplementary Figure S4.** TMZ-induced mutations in the collection of TK6 DNA repair mutants. **(A)** Box plots indicating statistical significance of differences in mutation numbers among the cell lines in Figure 7C. Significant differences are indicated by # and * (# p value ≤0.1, * p value ≤ 0.05, ** p value ≤ 0.01, *** p value ≤ 0.001, t-test). **(B)** Numbers of indels (upper panel) and SVs (lower panel) induced per microM TMZ after subtracting untreated background. The dashed line corresponds to an average of WT. The statistical tests are in comparison to the WT line. **(C)** Chromatin fractionation assay. Cells were treated with LD_90_ concentration of TMZ (1.25 microM for *ATAD5-/-*, 5 microM for WT) for 24 hours. **(D)** Numbers of indels induced per microM TMZ after subtraction of untreated background in *MSH2-/- ATAD5-/-* line. **(E)** Numbers of dinucleotide deletions in poly A repeats longer than 12 nt induced in the POLB-*/-* line per microM TMZ after subtraction of untreated background.
